# Uncovering Design and Assembly Rules for mRNA–DNA Origami

**DOI:** 10.1101/2025.10.09.681283

**Authors:** Jack Y. Wang, Jared Huzar, Myoungseok Kim, Taeyoung Ryu, Nahtalee R. Lomeli, Derfogail Delcassian, Do-Nyun Kim, Grigory Tikhomirov

## Abstract

mRNA–DNA hybrid origami offers a powerful route to combine the structural programmability of DNA origami with the biological functionality of messenger RNA, but generalizable design and assembly rules for these hybrids remain poorly defined. Here we systematically investigate the design principles and synthesis conditions that govern high-yield formation of mRNA–DNA hybrid nanostructures. Using mature mRNAs encoding firefly luciferase, EGFP, and mCherry as scaffolds, we construct a series of five hybrid compact origamis with diverse sizes, shapes, crossover strategies, and packing densities. We identify key parameters that control folding fidelity, including asymmetric A-form crossovers, monovalent-cation concentrations, and moderate-temperature annealing protocols, which together mitigate RNA instability, reduce kinetic traps, and accommodate RNA–DNA helical geometry. Atomic force microscopy reveals monodisperse, well-folded structures consistent with design expectations across most architectures and confirms that optimized conditions produce nanoscale precision comparable to DNA origami. Our findings establish generalizable design rules and a standard synthesis protocol for mRNA–DNA hybrid origami, providing a framework for their use in gene delivery and other RNA-based nanotechnologies.

---

Nucleic-acid origami has transformed molecular nanotechnology by enabling the folding of a long single strand into complex three-dimensional shapes via complementary oligonucleotide staples^1,2^. The well-defined design principles and synthesis conditions of DNA origami enabled the construction of nano-scale robotics^3–7^, precision measurement systems^8^, and biosensors^9–12^. While DNA origami has matured into a powerful and versatile platform, the integration of RNA into these architectures holds unique promise: RNA can act not only as a structural scaffold but also as a functional biomolecule, carrying coding regions, aptamers, ribozymes, or regulatory elements^13–15^. Blending the programmability of DNA with the biological functionality of RNA could enable next-generation nanostructures for gene delivery, synthetic biology, and responsive therapeutics.

Early efforts in RNA origami focused on motif-based systems and modular units (e.g. tectoRNAs) that self-assemble via kissing loops or fixed junctions^16^. With advances in computational design tools and folding control, more recent strategies support co-transcriptional assembly, where scaffold and staple transcripts fold directly in one pot during in vitro transcription^17–19^. Such designs have enabled modestly sized shapes and functionalities, but they often rely on clever motif engineering and operate under narrow folding conditions.

Parallel to purely RNA strategies, RNA–DNA hybrid origami uses a long RNA scaffold folded by DNA staples, leveraging the rich design ecosystem of DNA while retaining RNA’s biological roles. Foundational works demonstrated that hybrid constructs can fold with reasonable yields and offer enhanced stability relative to simple duplexes^20–26^. For example, a “mini DNA–RNA hybrid origami nanobrick” used a 401-nt RNA scaffold folded by 12 DNA staples into a ten-helix bundle and showed greater resistance to RNase H digestion than duplex controls^22^. Another work developed design principles and a top-down algorithm to fold long RNA scaffolds with DNA staples into 3D wireframe polyhedra. Using mRNA, De Bruijn, M13 transcript, and 23S rRNA scaffolds, the authors build tetrahedra, octahedra, and pentagonal bipyramids, validate them by cryo-EM and DMS-MaPseq, and uncover base-level stability rules (e.g., GC placement near crossovers/termini) that guide sequence design and enable functional RNA scaffolds for delivery and structural studies^25^. These proofs of concept confirm that hybrid origami is feasible and potentially useful in biomedicine.

Among functional RNAs, messenger RNA (mRNA) has emerged as a leading therapeutic and functional biomolecule, spanning applications from simple cell-line transfection in basic research to organism-level immunomodulation and vaccination^27^. Its success hinges on efficient intracellular delivery and expression—parameters that are difficult to fine-tune with conventional carriers such as lipid nanoparticles (LNPs) or adeno-associated viruses (AAVs). LNPs often exhibit broad particle-size distributions and limited cell-type specificity, while AAVs are constrained by small genetic payloads and potential immunogenicity^27^. In contrast, DNA origami has been shown to offer programmable control over size, shape, and responsiveness to stimuli, as well as the ability to target specific cell types and anatomical sites, making it a promising delivery vehicle for therapeutic cargo^28^. Incorporating mRNA as the scaffold in nucleic-acid origami could therefore deliver unprecedented programmability and versatility to gene-delivery systems, potentially overcoming current limitations in targeting specificity and chemical modification^27^.

However, multiple barriers must be addressed before mRNA-scaffolded origami can become a viable therapeutic platform. First, the field lacks robust design principles for RNA–DNA hybrid origami, making high-yield assembly of diverse structures challenging. Second, several intrinsic properties of RNA prevent direct transfer of DNA-origami rules to mRNA–DNA hybrids. For example, the C2 ′-hydroxyl group in RNA renders it susceptible to autolysis at elevated temperatures^29,30^ and to degradation at high divalent-cation concentrations^31,32^ — conditions that can compromise downstream gene expression. Third, RNA is more prone than DNA to “kinetic traps,” misfolding into undesired secondary structures during annealing^33,34^. Finally, conformational differences between A-form RNA and B-form DNA complicate direct translation of crossover placement, helical parameters, and other design features from DNA to RNA or RNA–DNA hybrids.

To fully exploit mRNA-scaffolded origami as a next-generation platform for molecular therapeutics and gene delivery, systematic studies are needed to elucidate its design principles, folding pathways, and synthesis conditions as done herein. We expect that this work and others will lay the foundation for reproducible, high-fidelity hybrid nanostructures that combine the structural programmability of DNA with the functional potential of RNA.

## Results and Discussion

We synthesized a series of compact mRNA–DNA hybrid origami nanostructures and identified key design and assembly parameters that enable high-yield folding (Figure 1). These hybrids integrate the structural programmability of DNA origami with the functional potential of mRNA. Formation of each nanostructure relied on pre-programmed base pairing between an mRNA scaffold and complementary single-stranded DNA staples, facilitated by cations and heat (Figure 1a)^2,25^. Building on the established scaffold–staple model of DNA origami, we adapted crossover placement, helix parameters, and annealing conditions to accommodate the distinct geometry and chemistry of RNA.

**Figure 1.**
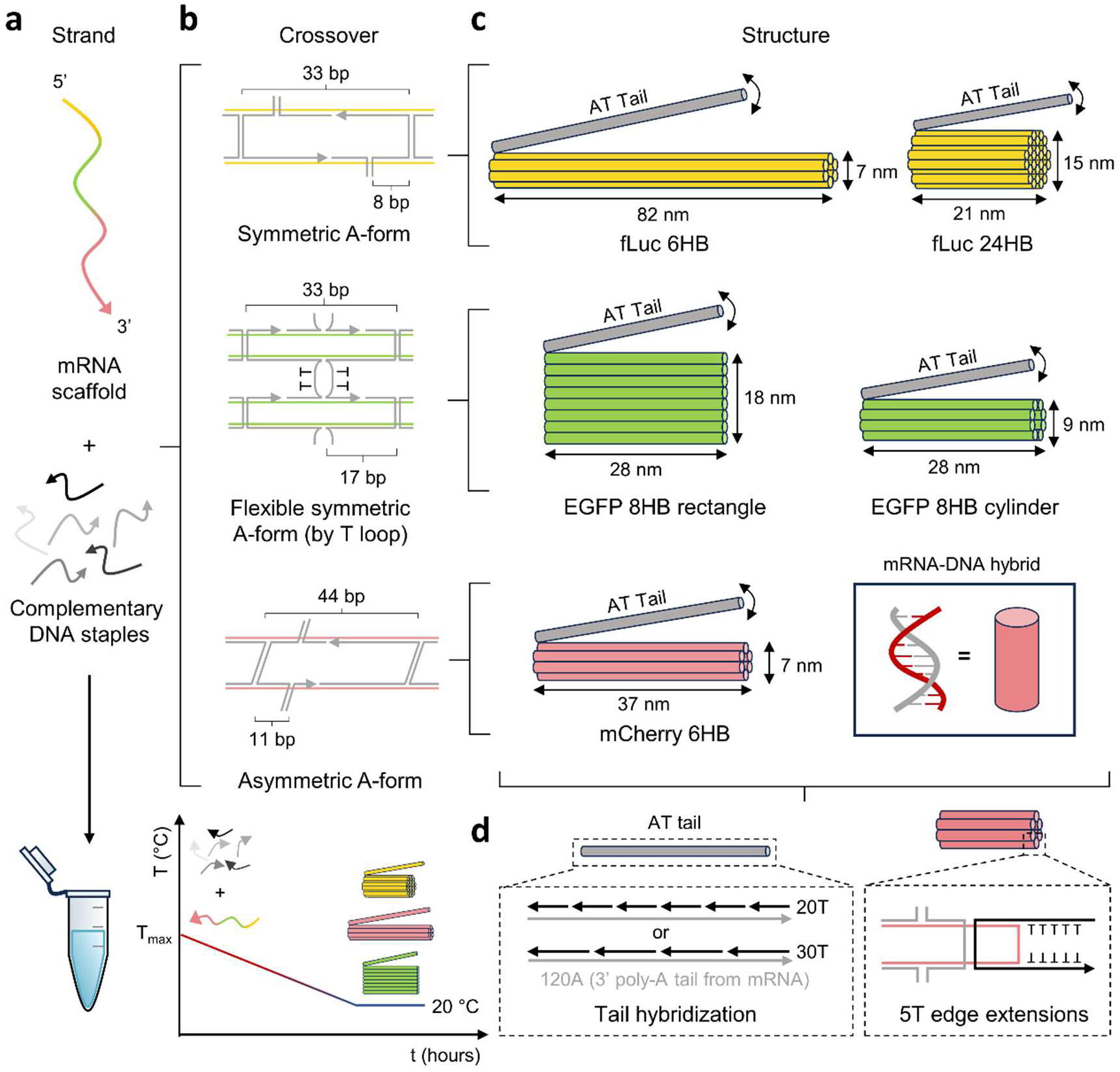
mRNA-DNA hybrid origami design. (a) Self-assembly of mRNA-DNA hybrid origami through thermal incubation of mRNA scaffold and complementary DNA staples. (b) Double crossover designs implemented in each origami nanostructure. fLuc, EGFP, and mCherry-encoding mRNA scaffold structures use symmetric A-form crossovers, flexible symmetric A-form crossovers, and asymmetric A-form crossovers, respectively. (c) Schematics dimensions for each origami calculated with A-form parameters. Each cylinder represents a duplex helix bundle. (d) Common features of mRNA-DNA hybrid origami, including 3’ AT tail and 5T edge staple extensions.

To establish a generalizable protocol for mRNA–DNA origami synthesis, we designed and folded five diverse nanostructures encoding different proteins. Specifically, we used mature mRNAs encoding firefly luciferase (fLuc), enhanced green fluorescent protein (EGFP), and mCherry as scaffolds for an array of hybrid origami architectures (Figure S1a-e).

To assess how design parameters influence the structural stability of mRNA–DNA hybrid origami, we designed, synthesized, and characterized five nanostructures with systematically varied properties. We focused on parameters known to shape structural and functional performance: (i) aspect ratio and (ii) dimensionality, which affect cellular uptake and biodistribution^28^, (iii) crossover pattern design, which governs stability under physiological conditions^35–37^, (iv) scaffold length, which determines gene-encoding capacity, and (v/vi) helix arrangements and packing density, which together control gene-loading density for potential therapeutic applications.

### Crossover pattern design

We first isolated the effect of staple crossover geometry on origami conformation to test whether DNA-origami crossovers can be applied directly to mRNA–DNA hybrids (Figure 1b). All structures were designed for an A-form helix (11 bp per turn) in line with established RNA-origami rules^20,22,25^. We evaluated whether relatively sparse A-form crossover densities (33 bp for fLuc and EGFP, 44 bp for mCherry) could still produce stable origami compared with the denser 21–32 bp crossovers typical of DNA origami. We also contrasted perpendicular double crossovers (fLuc) with diagonal crossovers (mCherry) to test the importance of tailoring crossovers specifically to A-form geometry. The fLuc structures used symmetric, perpendicular A-form double crossovers adapted from conventional DNA origami, whereas the mCherry 6HB employed asymmetric, diagonal crossovers adapted from RNA origami (Figure 1b–c). Although perpendicular crossovers are common in DNA origami, they do not account for the steeper axial inclination of hybrid RNA-DNA duplexes relative to B-form DNA^25,38,39^ and can induce torsion and distortion. To mitigate this, we modified A-form double crossovers with T-loop inserts in the EGFP origami to enhance interhelix flexibility^40^ and compensate for potential internal torsion. Together, these designs allowed us to test the feasibility and precision of distinct double-crossover strategies in mRNA–DNA hybrid origami.

### Common structural features

Despite differences in crossover pattern, all hybrids incorporated design elements to manage the repetitive scaffold 3′ poly(A) tail and reduce blunt-end aggregation. Each structure included a 120 bp AT “tail,” formed by hybridizing the 3′ poly(A) segment to four 30 nt poly(T) oligos (fLuc 6HB, fLuc 24HB) or six 20 nt poly(T) oligos (EGFP 8HB rectangle, EGFP 8HB cylinder, mCherry 6HB, Figure 1d; Figure S1a-e). Additionally, all edge staples carried 5T single-stranded extensions to minimize blunt-end stacking and interparticle aggregation^41^ (Figure 1d; Figure S1a–e).

### Computational validation

To predict and verify our design principles, we performed molecular-dynamics simulations of each origami in strict A-form, strict B-form, and intermediate conformations using SNUPI^42^ and oxDNA^43^. Static SNUPI analyses^42^ revealed that all hybrid origami formed overall rigid structures with only modest flexibility at helix-bundle edges comparable to DNA origami (Figure 2a). Notably, simulations exposed marked differences between A-form, B-form, and hybrid geometries, underscoring the impact of crossover design on structural integrity. For example, the fLuc origami’s equilibrium structure aligned best with the intended design under B-form simulation, but showed poorer agreement under A-form or hybrid simulations. These results indicate that perpendicular crossovers introduce internal torsion in RNA-DNA hybrids. This effect was evident in the fLuc 6HB, where two “joints” appeared in the SNUPI hybrid and oxDNA A-form simulations but not in the oxDNA B-form simulation (Figure 2b), indicating torsional distortion from orthogonal double crossovers. Although adding interhelix flexibility reduced this distortion, it did not fully restore the structural integrity achievable in DNA–DNA origami, as reflected by inward folding of the EGFP 8HB. In contrast, the mCherry 6HB, designed with asymmetrical diagonal crossovers, exhibited minimal structural distortion, highlighting the advantage of diagonal crossovers for accommodating the greater axial inclination of RNA–DNA duplexes.

**Figure 2.**
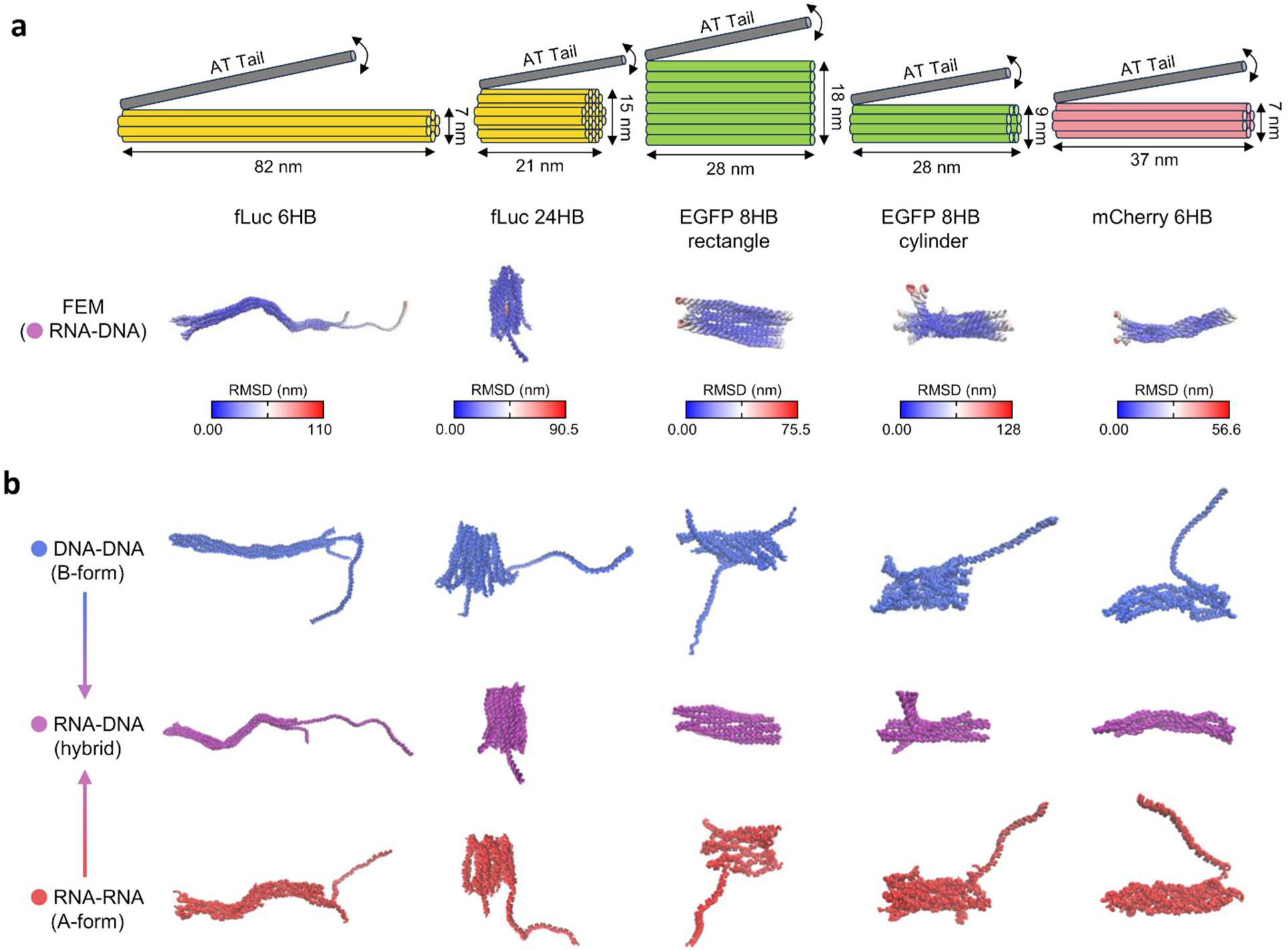
Molecular dynamics simulations of hybrid origami structures. (a) Hybrid origami parameters from A-form DNA calculations and corresponding root-mean-square deviation (RMSD) with finite element method representations of each origami structure. Simulations were performed using Structured NUcleic acids Programming Interface (SNUPI) with RNA-DNA hybrid conformation. (b) Origami equilibrium structure simulations with DNA-DNA (B-form, oxDNA), RNA-DNA (hybrid, SNUPI), and RNA-RNA (A-form, oxDNA) parameters, respectively. All structures excluding fLuc 6HB were omitted of AT tails for SNUPI simulations.

Taken together, these findings emphasize that RNA–DNA helices possess unique geometric constraints that must be explicitly considered in hybrid origami design. Developing refined crossover strategies tailored to A-form geometry is essential for achieving high-fidelity, torsion-free mRNA–DNA hybrid nanostructures.

### Synthesis parameter optimization

Having defined how crossover geometry and nucleic acid conformation influence hybrid origami structure in silico, we next examined how synthesis conditions should be adapted from conventional DNA-origami protocols. We began by testing the role of cation species and concentration, as monovalent and divalent cations including Na⁺ and Mg^2+^ reduce electrostatic repulsion and are widely used to promote DNA-origami folding^44–46^ (Figure 3a). High MgCl_2_ concentrations, however, may be unsuitable for mRNA scaffolds because RNA’s A-form geometry and C2 ′-hydroxyl group confer sensitivity to autolysis at elevated divalent-cation concentrations^31,32^. To identify optimal conditions, we synthesized our mRNA–DNA hybrids in varying concentrations of NaCl and MgCl_2_. As an initial screen, we tested whether hybrid origami could form at Mg²⁺ levels typical for DNA origami (5–20 mM). Agarose gel electrophoresis (AGE) revealed faint structural bands for fLuc 6HB and fLuc 24HB at 6–10 mM MgCl_2_, whereas mCherry 6HB and EGFP 8HB rectangle displayed no detectable bands at 10–20 mM MgCl_2_ (Figure 3b), indicating that DNA-origami protocols are not directly transferable to mRNA–DNA hybrids. We hypothesized that eliminating divalent cations could improve folding yield by enhancing RNA stability^25^. Indeed, origami assembled in NaCl as low as 60 mM exhibited stronger structure bands under otherwise identical buffer conditions (Figure 3c). Based on its higher yields and broader effective range, NaCl was selected as the standard cation for mRNA–DNA hybrid origami assembly.

**Figure 3.**
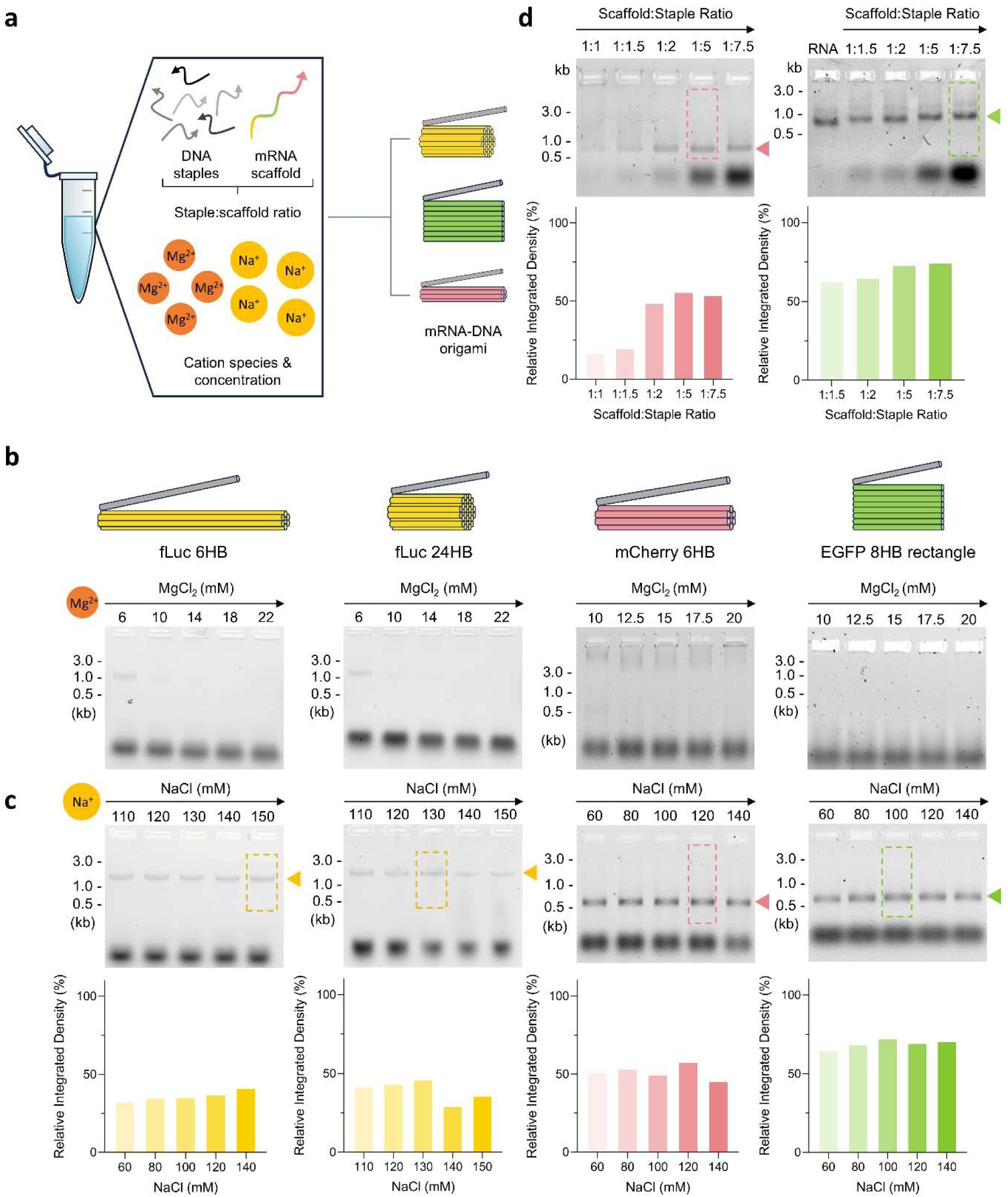
Buffer condition optimization for mRNA-DNA hybrid origami synthesis. (a) Overview of synthesis buffer components. Cationic species are required for facilitating correct folding of staples and scaffold into origami. (b) Synthesis of mRNA-DNA origami using varying concentrations of MgCl_2_ and relative structure band intensity. (c) Synthesis of hybrid origami using varying concentrations of NaCl and relative structure band intensity. (d) Scaffold:staple molar ratio screening for optimal formation of mCherry 6HB and EGFP 8HB rectangle. DNA staples were added in 1 to 7.5 molar equivalents to 10 nM of mRNA scaffold. Agarose gel electrophoresis (AGE) results (top) and relative band intensity of structure bands (bottom) are shown. All origami were synthesized using the same 4-hour annealing protocol. Structure band positions are indicated by triangles, lanes indicated by dashed boxes. AGE was performed with 1% agarose by mass in 1× sodium-borate buffer, stained with 8 μL/mL SYBR Safe, run at 200 V for 35 minutes. Ladder size bands are indicated on the left of gels, derived from DNA ladder lanes (not shown). RNA lanes contain 10 nM of RNA.

We next performed a NaCl concentration screen to define a generalized optimal range for hybrid origami folding. AGE band-intensity analysis indicated robust yields for fLuc 6HB, fLuc 24HB, mCherry 6HB, and EGFP 8HB rectangle between 60 and 140 mM NaCl (Figure 3c). Notably, NaCl concentrations as low as 60 mM supported origami formation, whereas optimal folding for most mRNA–DNA hybrids occurred at 110–120 mM. Accordingly, we adopted 115 mM NaCl as our standard assembly condition.

We then optimized the scaffold-to-staple molar ratio, anticipating that the ideal ratio would differ from DNA origami because RNA–DNA duplexes have different thermodynamic stabilities than DNA–DNA duplexes^47^. AGE analysis revealed maximal mCherry yields at ratios between 1:2 and 1:7.5, with comparable trends for EGFP 8HB rectangle (Figure 3d). Remarkably, ratios as low as 1:1.5 were sufficient for origami formation in both structures, consistent with reports that RNA–DNA origami can require far less staple excess than DNA origami^2,22,25^. We attribute this greater staple efficiency to stronger base stacking and 2′-OH stabilization in RNA^48^. For consistency across designs, we standardized a 1:7.5 scaffold-to-staple ratio for subsequent experiments.

Finally, we assessed how the annealing protocol influences hybrid-origami yield. Thermal incubation drives DNA-origami assembly and prevents kinetic traps^2^, but typical annealing ramps may be incompatible with RNA–DNA hybrids because RNA is both more prone to kinetic traps and more susceptible to autocatalysis at elevated temperatures^33,34^. Temperatures that are too low may fail to disrupt RNA secondary structures and thus reduce yield, whereas too high can degrade RNA. We first tested an isothermal protocol originally developed for rapid DNA-origami assembly at physiological conditions^49^ and compared it with a standard temperature-ramped annealing protocol. Specifically, we evaluated 4-h isothermal versus 4-h annealing conditions for both EGFP 8HB rectangle and EGFP 8HB cylinder to determine whether the effects are different for 2D versus 3D architectures, as observed for some DNA origami^2^ (Figure 4a–b). AGE analysis showed comparable yields between 2D and 3D structures within each protocol but consistently higher yields for the annealing ramp versus the isothermal protocol at the same temperature (Figure 4c). These results indicate that isothermal incubation at 55 °C can assemble mRNA–DNA hybrid origami but that annealing ramps at the same temperature produce superior yields. Further optimization of the maximum temperature for isothermal protocols may enhance efficiency and suppress kinetically favored byproducts.

**Figure 4.**
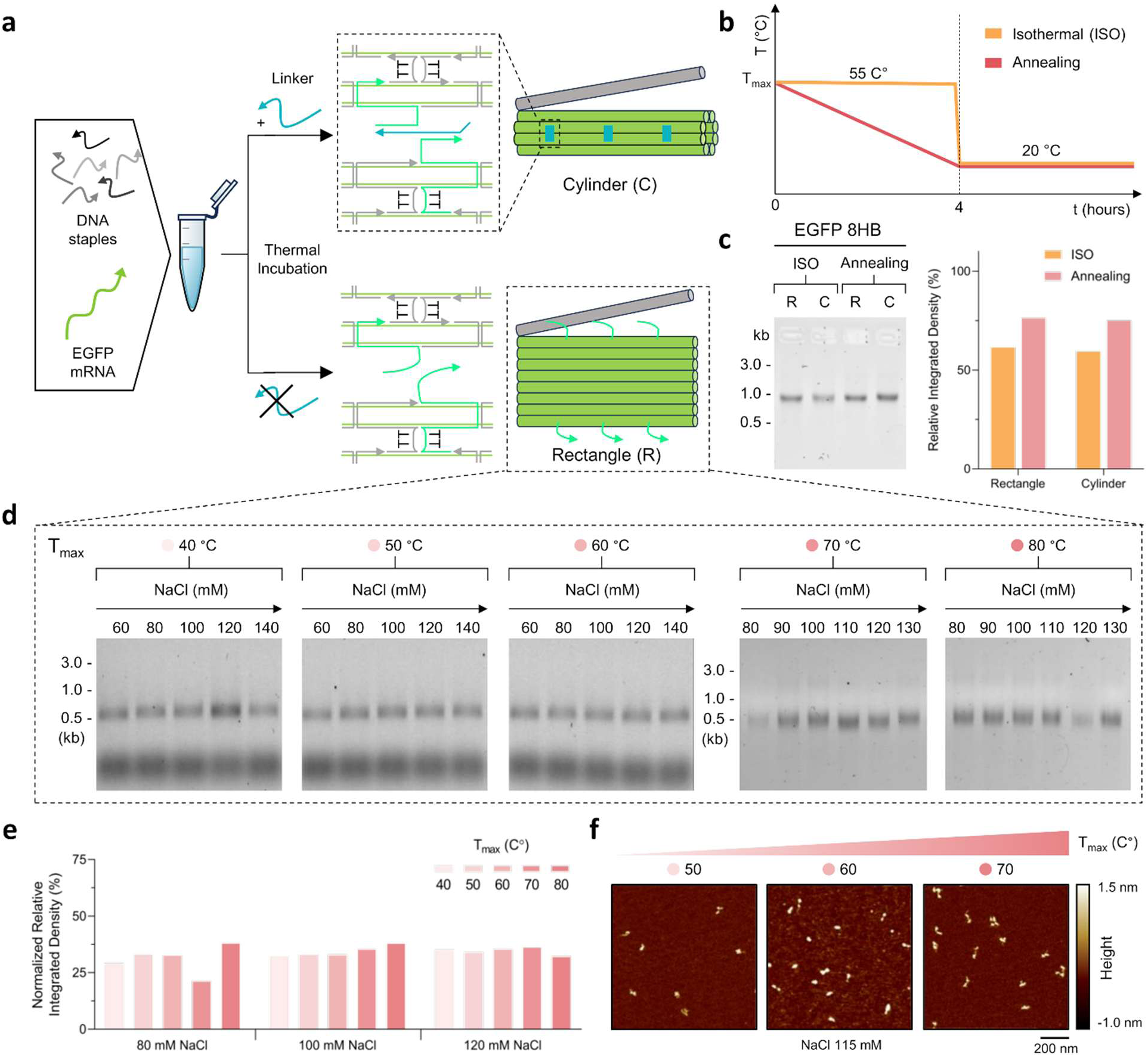
Incubation protocol optimization for mRNA-DNA hybrid origami synthesis. (a) Addition of linker strands (cyan) facilitates formation of EGFP 8HB cylinder rather than EGFP 8HB rectangle. (b) Schematic of 4-hour isothermal (ISO) and annealing protocols. Vertical axis indicates temperature in °C, horizontal axis indicates time in hours. (c) AGE results (left) and structure band intensities (right) for EGFP 8HB rectangle (R) and cylinder (C) synthesized with ISO and annealing protocols, respectively. (d) AGE results (top) and structure band intensity (bottom) of EGFP 8HB rectangle synthesized at different T_max_ and NaCl concentrations. AGE was performed with 1% agarose by mass in 1× SB buffer, stained with 8 μL/mL SYBR Safe, run at 200 V for 35 minutes. DNA size is shown on the left of gels, derived from DNA ladder (not shown). Staple bands are excluded from (c) and T_max_ = 70 and 80 °C gels due to scaling.

After establishing the optimal incubation protocol, we next determined the maximum temperature (T_max_) that yields the most efficient folding. We performed a comprehensive screen of EGFP 8HB rectangle origami assembled at different T_max_ values (each with matched ramp rates) and NaCl concentrations using 4-hour annealing protocols (Table S6). AGE images and corresponding band-intensity analysis showed comparable structural bands across T_max_ values from 40 to 80 °C and NaCl concentrations between 80 and 140 mM (Figure 4d). These data indicate that temperatures in the range of 40–80 °C combined with 60–140 mM NaCl are sufficient to support proper folding of mRNA–DNA hybrid origami (Figure 4d).

Notably, band intensities were most consistent at T_max_ values of 50–60 °C across all sodium concentrations tested, suggesting a generally optimal temperature window (Figure 4e). AFM analysis further corroborated these findings: images of EGFP 8HB rectangle structures synthesized at T_max_ = 60 °C were more uniform in shape and contained fewer partially folded intermediates compared to those assembled at T_max_ = 50 °C or 70 °C (Figure 4f). We infer that temperatures above 70 °C may accelerate RNA autocatalysis and hinder folding yield, whereas T_max_ below 50 °C may leave RNA secondary structures intact and reduce staples incorporation. Based on these results, we adopted 55 °C as the standard T_max_ for synthesis of EGFP-derived mRNA–DNA hybrid origami and generalized this condition to other designs to maximize both structural integrity and yield.

### AFM characterization

To further characterize and verify the formation of our mRNA–DNA hybrid origami, we performed atomic force microscopy (AFM) imaging of each structure. AFM reconstructions revealed monodisperse, well-folded architectures for all origami synthesized under our standard conditions (115 mM NaCl; T_max_ = 55 °C, 4-hour annealing), with the exception of the EGFP 8HB cylinder, thereby confirming the general applicability of these synthesis parameters (Figure 5a). Additional AFM imaging of the EGFP 8HB cylinder showed a mixture of completely and partially folded species, indicating strong interaction with mica surface (Figure S3a–c).

**Figure 5.**
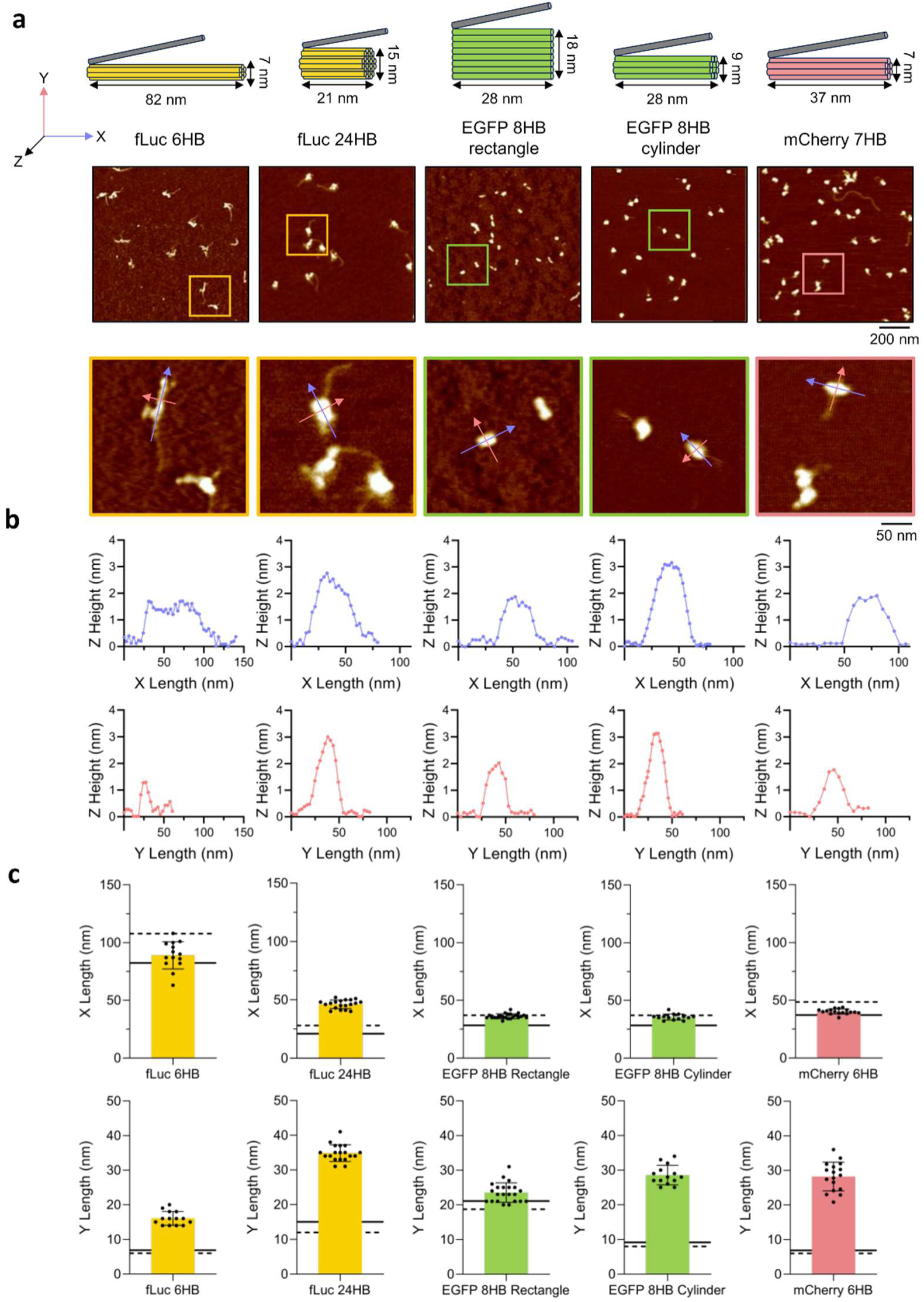
AFM characterization of mRNA-DNA origami. (a) 1×1 μm (top) and zoomed in 250×250 nm (bottom) air AFM images of each hybrid origami nanostructure. X, Y and Z axes were defined by helical axial length, helical bundle width, and height, respectively. Pink and blue arrows in 250×250 nm air AFM images correspond to X and Y axes, respectively, for section analysis. Origami measurements were derived from A-form conformation parameters. (b) Origami section data for X and Y axes derived from labeled structures in 250×250 nm air AFM images. Vertical axis corresponds to Z axis height, with horizontal axes corresponding to X axis (top) and Y axis lengths (bottom). (c) Mean ± SD of X (top), Y (middle) and Z (bottom) lengths for each origami. Sections were sampled from individual origami in 1×1 μm air AFM images, with sample size of *N* = 14, 19, 17, 22, 14 for fLuc 6HB, fLuc 24HB, mCherry 6HB, EGFP 8HB rectangle and EGFP 8HB cylinder, respectively. A-form and B-form estimates for each structures’ axis lengths are indicated with black-colored solid and dashed lines, respectively.

Higher-resolution air-AFM reconstructions showed features consistent with design expectations: most individual origami displayed a flexible extension corresponding to the AT tail (absent only in the EGFP 8HB rectangle) and a dense region matching the helix bundles, further confirming proper folding (Figure 5a).

After establishing qualitative resemblance to design features, we quantified origami fidelity by comparing experimental dimensions to theoretical predictions. We measured axial length (x) and helical width (y) across RNA–DNA helices. Mean axial lengths fell between estimates from strict A- and B-form conformations for all designs except fLuc 24HB (Figure 5b–c), consistent with reports that RNA–DNA duplexes adopt an intermediate conformation between A and B forms^50–52^. In contrast, mean helical widths were two- to threefold greater than estimated for all structures except EGFP 8HB rectangle (Figure 5b–c). We attribute this discrepancy to imaging artifacts inherent to air AFM, including tip-convolution effects^53^.

To verify AFM tip effects, we imaged a similar DNA 6HB control (predicted helical length 337 nm, width 7 nm) under the same conditions (Figure S4a). AFM reconstructions of this DNA 6HB mirrored the hybrid-origami trend: measured lengths matched estimates while measured widths were two- to threefold larger (Figure S4d). We therefore conclude that width discrepancies reflect AFM imaging bias rather than the lack of structural integrity.

Collectively, these AFM characterizations confirm both the structural fidelity of mRNA–DNA hybrid origami and the robustness of the design and synthesis parameters across diverse nanostructures with varying sizes, bundle geometries, aspect ratios, and crossover densities. Together, these findings demonstrate that our optimized conditions and design rules are generalizable across a wide range of RNA–DNA hybrid origami architectures.

## Conclusion

We have constructed and systematically studied a diverse set of compact mRNA–DNA hybrid origami nanostructures spanning distinct sizes, shapes, staple crossover strategies, packing densities, and gene-encoding scaffolds. By dissecting the impact of individual design features and synthesis conditions, we have established generalizable parameters for the high-yield assembly of hybrid origami. Across all designs tested, we found that asymmetric A-form crossovers, elevated monovalent cation concentrations, and moderate-temperature annealing protocols consistently improved folding yield and structural fidelity. These improvements likely reflect mitigation of RNA instability, reduction of kinetic traps, and explicit accommodation of RNA–DNA helical geometry. On the basis of these findings, we propose a standardized synthesis protocol that can serve as a starting point for the formation of mRNA–DNA hybrid origami for gene delivery and other applications.

While this study provides a foundation for reliable hybrid-origami assembly, further work is needed to refine synthesis, characterization, and in situ performance. In particular, the stability of individual structural motifs under biologically relevant conditions and their impact on mRNA translation efficiency merit systematic investigation. Such studies will be essential to fully unlock the potential of hybrid origami for therapeutic and diagnostic use.

Despite these remaining challenges, the present work establishes a framework of design rules and synthesis conditions that makes mRNA–DNA hybrid origami more predictable and accessible. Leveraging the high programmability and versatility of this platform could open new avenues in precision medicine, gene delivery, and nano-therapeutics, accelerating translation of hybrid origami from proof-of-concept to practical biomedical tools.

## Methods

### Scaffold and staple design

All mRNA-DNA hybrid origami nanostructures were designed with browser-based design tool ScadNano^54^. Complete mCherry, fLuc and EGFP scaffold mRNA sequences (including open reading frame, 5’ and 3’ untranslated regions (UTRs)) were provided by TriLink BioTechnologies. Origami dimension parameters were calculated using A-form DNA parameters of 11 bp per turn, 2.55 Å axial rise per bp, and 23 Å helix diameter. fLuc 6HB, fLuc 24HB, EGFP 8HB rectangle, EGFP 8HB cylinder, and mCherry 6HB each consist of mRNA scaffold encoding for the structure’s corresponding fluorescent protein and 57, 56, 35, 39 and 30 complementary DNA staple oligos, respectively (Figure S1a-e, Table S1-S5). A single nucleotide mismatch was induced at position 1 of the scaffold 5’ sequence to facilitate strand displacement at the 5’ UTR for future studies of gene delivery.

### Molecular dynamics simulations

For oxDNA simulations, each origami nanostructure was simulated to acquire equilibrium configuration, root-mean square fluctuations (RMSF), and energy distribution histograms. Simulations of DNA-DNA and RNA-RNA interactions of each nanostructure were performed to predict an intermediate RNA-DNA hybrid structure. Structured NUcleic acids Programming Interface (SNUPI) was further used to acquire predictions of origami in equilibrium, in addition to origami structural properties and RMSF of structures without AT tails. The nonlinear properties of ssDNA and ssRNA were considered for the above SNUPI simulations^55–57^. The RMSF of structures were obtained by normal mode analysis (NMA). Step parameters used in the SNUPI simulations were obtained using all-atom molecular dynamics simulations.

### Cation concentration screening

fLuc 6HB, fLuc 24HB, EGFP 8HB rectangle and mCherry 6HB were synthesized in synthesis buffer at varying concentrations of NaCl or MgCl_2_. Synthesis buffer conditions include 10 mM Tris and 1× RNAsecure (Invitrogen) containing 10 nM mRNA scaffold and 75 nM of each staple, with variable cation concentrations of 6-22 mM MgCl_2_ or 60-140 mM NaCl at total volume of 100 μL per equivalent. Origami was synthesized using the same 50-20 °C 4-hour annealing protocol, then loaded and run in electrophoresis gels for analysis of structure band intensity.

### Staple:Scaffold ratio screening

EGFP 8HB rectangle and mCherry 6HB were synthesized using varying staple to scaffold ratios. 10-75 nM of each staple oligo were added to 10 nM of mRNA scaffold in total volume of 50 μL of 1× synthesis buffer (10 mM Tris base, 1× RNAsecure, 115 mM NaCl containing 10 nM mRNA scaffold and 75 nM of each staple oligo). Mixtures were then incubated using the same 50-20 °C annealing protocol for formation of origami nanostructures. Origami products were loaded and run in electrophoresis gels for analysis of band intensity to compare relative yield.

### Maximum annealing temperature screening

EGFP 8HB rectangle was synthesized at different initial T_max_ values using a constant temperature gradient (Table S6). Origami was synthesized in 50 μL of 1× synthesis buffer (10 mM Tris base, 1× RNAsecure, containing 10 nM mRNA scaffold and 75 nM of each staple oligo) at 60-140 mM NaCl, and were incubated at initial T_max_ = 40-80 °C to a final temperature of 20 °C for a total duration of 4 hours. Successfully synthesized origami samples were then collected and run in electrophoresis gels, then analyzed for band intensity to compare relative yield.

Incubation protocol screening: EGFP 8HB rectangle and EGFP 8HB cylinder were synthesized using isothermal or annealing protocols. Isothermal synthesis incubated origami synthesis mixtures at T_max_ = 55 °C for 4 hours, whereas annealing protocols incubated origami synthesis at initial T_max_ = 55 °C with continuous decrease to 20 °C for 4 hours with a consistent gradient. All origami nanostructures were formed in the same 1× synthesis buffer (10 mM Tris base, 1× RNAsecure, 115 mM NaCl containing 10 nM mRNA scaffold and 75 nM of each staple oligo). Nanostructures synthesized with both protocols were collected and run in electrophoresis gels for band intensity analysis of relative yield.

### Atomic force microscopy

2 μL of 10 nM origami were diluted 10× to a final volume of 20 μL with deposition buffer (15 mM MgCl_2_ in 1× TE buffer) and deposited on 10 mm diameter fresh mica slides (Bruker). Mica surfaces were incubated at room temperature for 5 minutes to allow nanostructure attachment, then washed with 200 μL ultrapure water for 3 times and blown dry with an air pump to remove excess nanostructures and salt crystals. Slides were loaded onto AFM (Bruker) and imaged with ScanAsyst-AirHR tip (Bruker) using NanoScope software and ScanAsyst Hi-Res program. Scan images were flattened to the 3rd order for sectional analysis with NanoScope Analysis software.

For sectional analysis, 1×1 μm and 250×250 nm air AFM images were flattened to the 3rd order using NanoScope Analysis software. Height sections for individual origami were taken for each origami nanostructure from corresponding 250×250 nm AFM images across the helical length (x axis), helical width (y axis), and height (z axis) using Section tool from NanoScope Analysis. Sections were further derived from 1×1 μm air AFM images to derive each dimensions’ length distribution for all origami. Sectional lengths were analyzed using GraphPad Prism 8 for mean and standard deviation.

### Agarose gel electrophoresis

Agarose gel electrophoresis (AGE) of EGFP-derived origami was performed with 150 mL of 1% agarose by mass (SeaKem) in 1× sodium borate (10 mM NaCl, 36.69 mM boric acid) buffer stained with 12 μL 1× SYBR-Safe (ThermoFisher Scientific) solution. AGE of fLuc and mCherry-derived structures were performed using the same conditions in 1× TBE buffer. Gels were solidified in gel cast with 15 or 20 well combs (BioRad). Sample wells contained 12 μL of nanostructure samples added with 3 μL of 6× loading dye (Invitrogen). Ladder wells contained 0.6 μL of 1 kb-plus DNA ladder (New England Biolabs) diluted with 11.4 μL of distilled water and 3 μL of 6× loading dye (Invitrogen). EGFP mRNA wells contained 0.322 μL of 3105.37 nM EGFP mRNA scaffold diluted with 11.7 μL distilled water and 3 μL of 10× gel loading dye (Invitrogen). Gel was run in an electrophoresis box (BioRad) with power supply at 200 volts (BioRad) for 35 minutes and imaged on UV tray with gel imager (BioRad).

For further gel band intensity analysis, AGE images were processed and analyzed using ImageJ. Structure bands were first identified, then gel lanes were defined using equal area boxes containing origami structure bands but excluding staple bands. Total integrated density of each gel lane box was acquired using ImageJ, with each relative band intensities calculated as percentages of structure band integrated density of total lane box integrated density.

## Supporting Information

The following files are available free of charge.

Additional design details, experimental protocols, materials and methods, including SNUPI simulation parameters and complete AGE images (PDF).

## Author Contributions

J.Y.W., J.H. and M.K. contributed equally to this work. J.Y.W., J.H. and M.K. designed, synthesized, and characterized mRNA-DNA nanostructures with help from N.R.L. T.R. modeled nanostructures in SNUPI. J.Y.W. wrote the manuscript with contributions from J.H., M.K., T.R., D.D., D.K., and G.T. G.T. guided the project.

The manuscript was reviewed by J.Y. Wang, J. Huzar, M. Kim, T. Ryu, N. R. Lomeli, and G. Tikhomirov. All authors have given approval to the final version of the manuscript. ǁ These authors contributed equally.

## Notes

The authors declare no competing financial interest.

## Acknowledgments

J.H. was supported by NIH training grant (T32GM146614). G.T. was supported by Society of Hellman Fellows Fund, UC Berkeley, NSF CAREER (2240000) and NSF POSE Phase 2 (2346048) awards. D.K. and T.R. were supported by the National Research Foundation of Korea (NRF) grant funded by the Ministry of Science and ICT (RS-2024-00346176).

## Abbreviations

mRNA: messenger RNA
ssDNA: single stranded
DNA bp: base pair
nt: nucleotide
AGE: agarose gel electrophoresis
AFM: atomic force microscopy
TEM: transmission electron microscopy

---

For Table of Contents Only

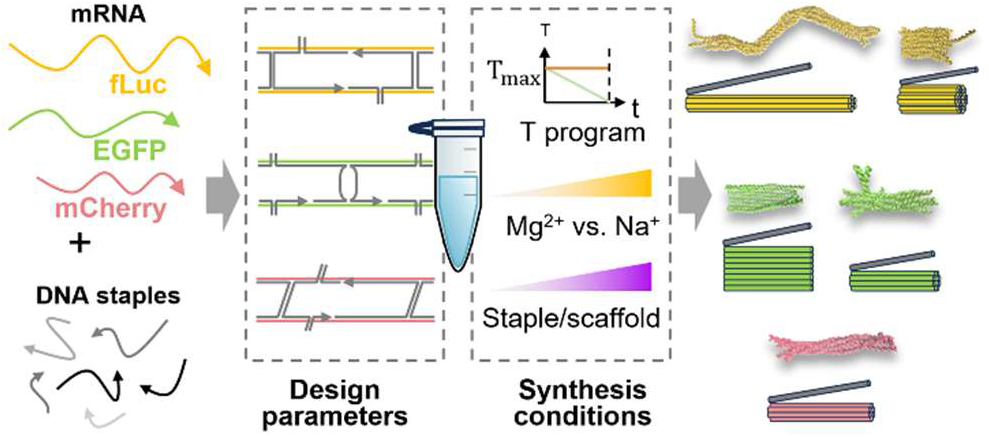

