## Supporting Information for "Uncovering Design and Assembly Rules for mRNA–DNA Origami"

### 1. Supporting Methods and Materials

**1.1 DNA 6HB design and synthesis**: We designed a DNA 6HB with well-defined parameters to serve as a control for AFM imaging. The DNA 6HB was designed using caDNAno software and consists of M13mp18 scaffold and 142 complementary staple oligos. Double crossovers spaced 21 bp apart were employed to connect adjacent 987 bp-long helix bundles and were arranged in a honeycomb lattice pattern. Excess M13mp18 scaffold was used to form ssDNA loop-outs in each helix bundle.

For synthesis of DNA 6HB, 100 nM of each staple oligo (IDT) from a 384-well plate were mixed using an acoustic liquid handler (Echo 525, Beckman) prior to addition to 7.5 nM M13mp18 scaffold (Bayou Biolabs) in 1×TE buffer (10 mM Tris and 1 mM EDTA, Fisher BioReagents) supplemented with 12 mM $MgCl_2$ (Invitrogen). The resulting mixture was incubated thermally using the following 12-hour protocol for origami formation: incubation at 90°C for 2 minutes, annealing from 90 °C to 86 °C (rate of -0.1 °C/second), 85 °C to 70 °C (rate of -0.1°C/30 seconds), 70°C to 40 °C (rate of -0.1°C/90 seconds), 40°C to 25°C (rate of -0.1°C/60 seconds), with final holding at 4°C.

Purification of DNA 6HB for AFM imaging was accomplished using the same protocol for mRNA-DNA hybrid origami described below, with 12 mM $MgCl_2$ in 1×TE buffer instead of 115 mM NaCl.

**1.2 mRNA-DNA hybrid origami purification**: To purify hybrid origami for use in proceeding stability assays, annealed origami products synthesized at standard conditions (10 mM Tris base, 1× RNAsecure, 115 mM NaCl containing 10 nM mRNA scaffold and 75 nM of each staple oligo) were purified using ultracentrifugation. 50 kDa filters (Amicon) were prewet with 500 µL of 115 mM NaCl and spun down in centrifuge tubes at 5000 rpm for 8 minutes. Filtrate was discarded prior to loading 50 µL of annealed origami and 450 µL of 115 mM NaCl and centrifuging for 5000 rpm for 8 minutes. The residual origami in the filter was washed three times. For each washing step, filtrate was discarded prior to addition of 450 µL of 115 mM NaCl to the residue and then spun down at 5000 rpm for 8 minutes. Residue containing purified origami was collected by inverting the filter in a new centrifuge tube and spun down at 10,000 rpm for 2 minutes.

**1.3 Quantification of mRNA-DNA hybrid origami concentration**: Nanodrop (Thermo Scientific) was used to determine hybrid origami concentration from ultracentrifugation purification, with each structure using a custom factor. Custom factor ($C_f$) calculations were based on linear interpolation of nucleic acid absorption coefficients: $C_f = \frac{33 \times n_{ss}}{(n_{ss}+n_{ds})} + \frac{50 \times n_{ds}}{(n_{ss}+n_{ds})}$, where $n_{ss}$ and $n_{ds}$ correspond to number of single-stranded and double-strand hybridized nucleotides. Nanodrop was blanked with 2 µL of 115 mM NaCl prior to performing measurements. Measurements were acquired using 2 µL of each structure with corresponding $C_f$ and concentrations of each origami were derived.

**1.4 mRNA-DNA hybrid origami degradation in 10% FBS-DMEM:** Purified mRNA-DNA origamis were incubated in 1× Dulbecco's Modified Eagle's Medium (DMEM, Gibco) supplemented with 10% FBS (VWR International) to assess origami stability in biological contexts. Ultracentrifugation-purified EGFP 8HB rectangle, EGFP 8HB cylinder, and mCherry 6HB were diluted with 10% FBS-DMEM to final concentrations of 10 nM at total volumes of 20 µL. The diluted origamis were then incubated at room temperature for 0, 1, and 10 minutes prior to loading into agarose gels for AGE with same conditions as previous.

**1.5 Additional SNUPI simulation information and parameters:** The intrinsic properties of mRNA-DNA hybrids were obtained using all-atom molecular dynamics (MD) simulations performed by NAMD with CHARMM36 force field for nucleic acids and TIP3P for water. RNA-DNA hybrid 6HB structures were solvated and ionized in a water box with 300 mM KCl concentration in a cubic periodic boundary cell with a padding distance of more than 15 Å. Timestep of 2 fs was used for the simulations. Short-range electrostatic potentials employed a cutoff of 12 Å and long-range electrostatic interactions were computed using particle-mesh-Ewald method with a grid spacing of 1 Å. The systems were initially subjected to energy minimization for 10,000 steps, followed up by thermalization of 0.2 ns. The equilibrium run was performed under 300 ns under the isobaric-isothermal (NPT) ensemble, where the pressure and temperature were maintained at 1 bar and 300 K using Nosé-Hoover Langevin piston scheme and Langevin thermostat, respectively. The intrinsic structural properties were calculated from the last 200 ns of the simulation trajectories.

From the molecular dynamics simulation trajectories, the step parameters were computed using the 3DNA definitions and were converted to suitable motif-dependent structural beam element properties. To analyze well-paired steps, base pairing was determined based on whether a hydrogen bond was formed or not between the N1 atom of a purine base and the N3 atom of a pyrimidine base with cut-off values of 4.0 Å and 40° for distance and angle, respectively. Additionally, for CO step and double CO step configurations, base pair steps with axial distances longer than 2.0 nm or shorter than 1.6 nm were excluded (average 1.8 nm).

#### 2. Scadnano Designs and HB Arrangements for Each Hybrid mRNA-DNA Origami

a

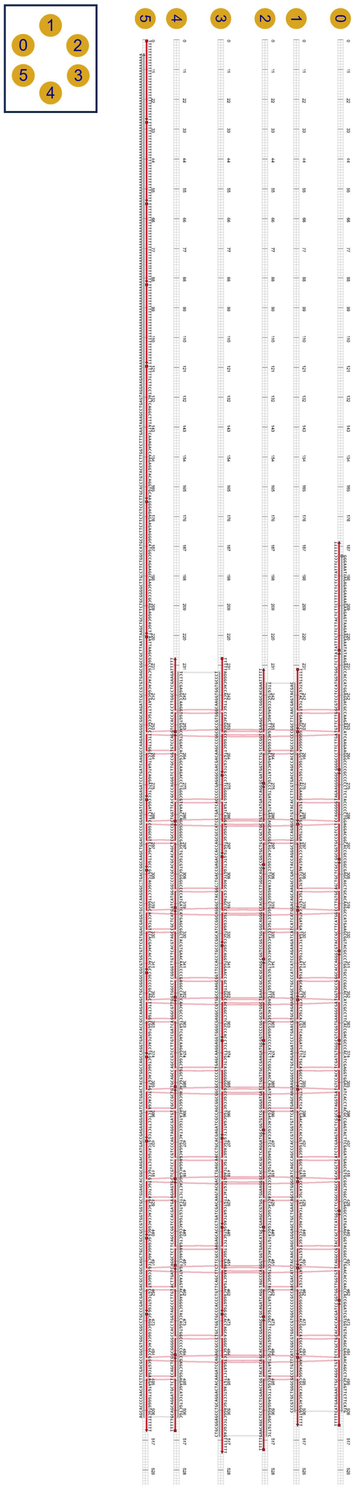

**b**

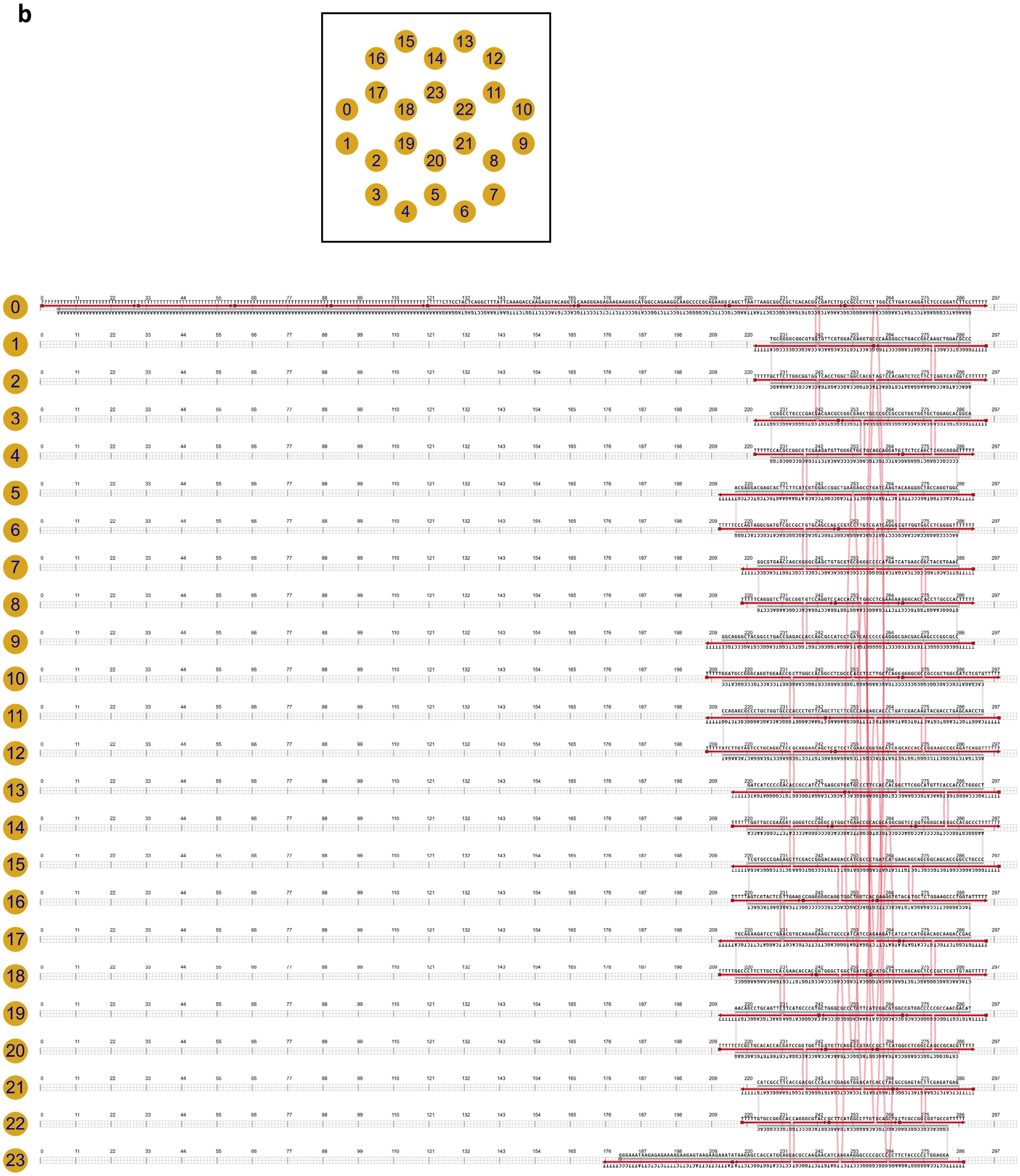
15
13
16
14
12
17
23
11
0
18
22
10
1
19
21
9
2
20
8
3
5
7
4
6

**c**

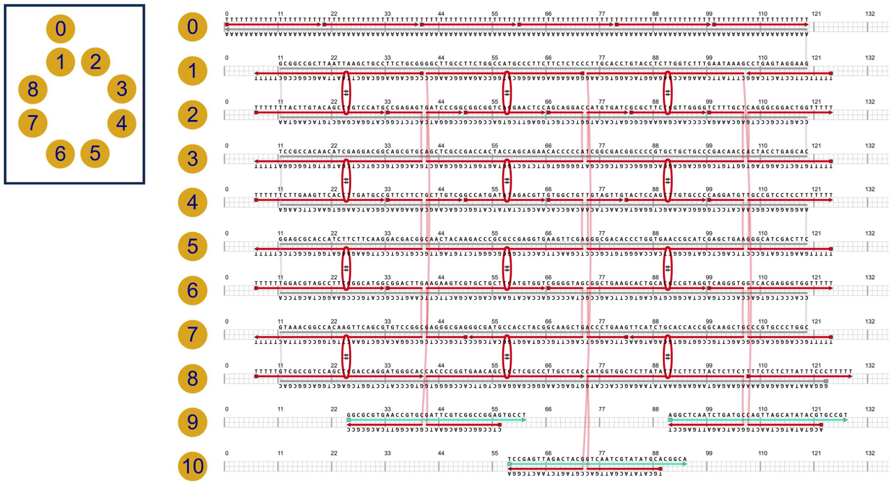

**d**

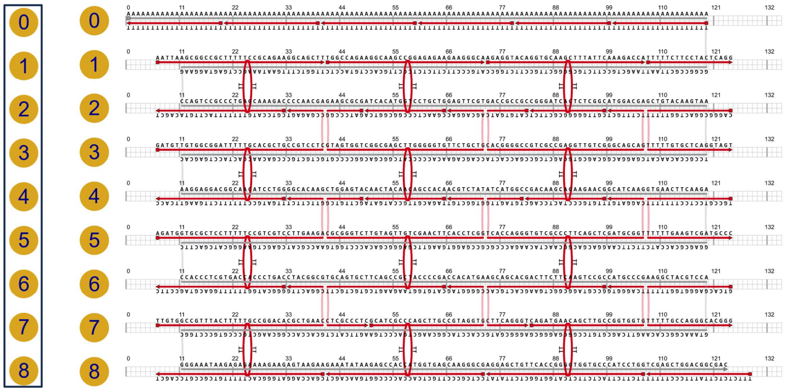

**e**

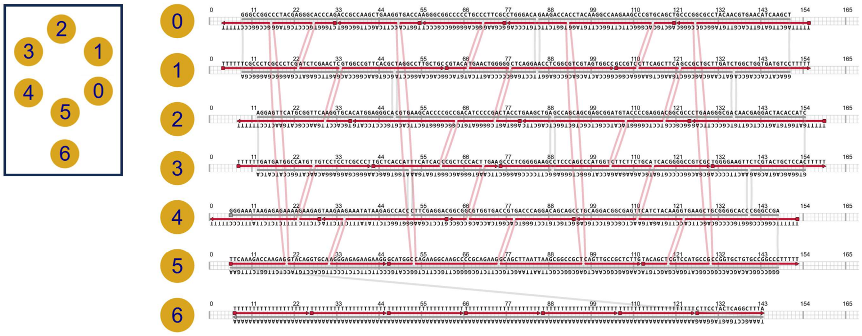

**Figure S1. Helix bundle side views and ScadNano designs for each origami.** (a) fLuc 6HB, (b) fLuc 24HB, (c) EGFP 8HB cylinder, (d) EGFP 8HB rectangle, and (e) mCherry 6HB. Helix numbers, scaffold (gray) sequence and staple (red) sequences are indicated.

### 3. Additional Molecular Dynamics Information for Each Origami

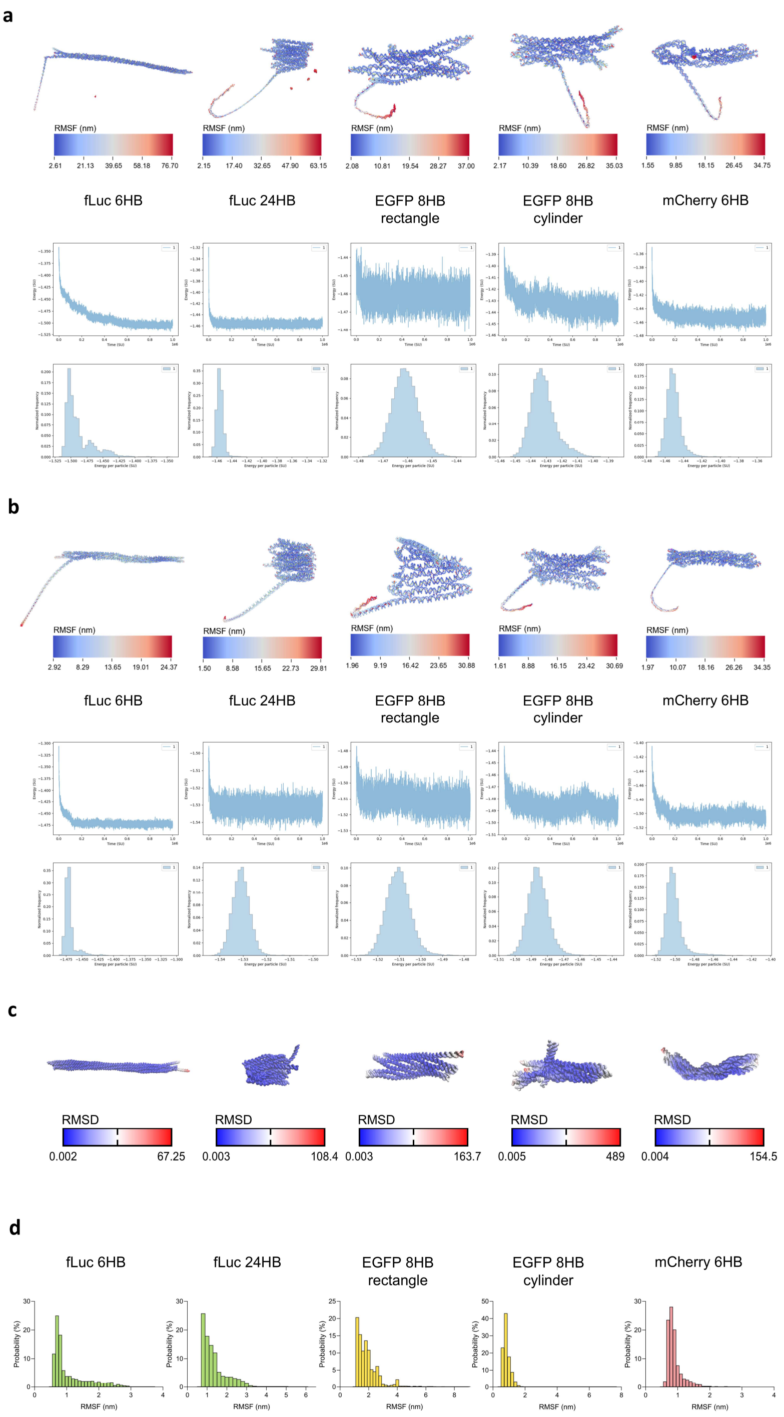

**Figure S2. Additional molecular dynamics simulation information for each origami**. oxDNA (a) DNA-DNA and (b) RNA-RNA simulations of each structures' RMSF, energy vs. time, and energy per particle distribution. (c) Additional strict B-form derived RMSD values from SNUPI for each structure. (d) RMSF probability density functions for each structure derived from RNA-DNA simulations using SNUPI.

#### 4. Additional AFM Sectional Data

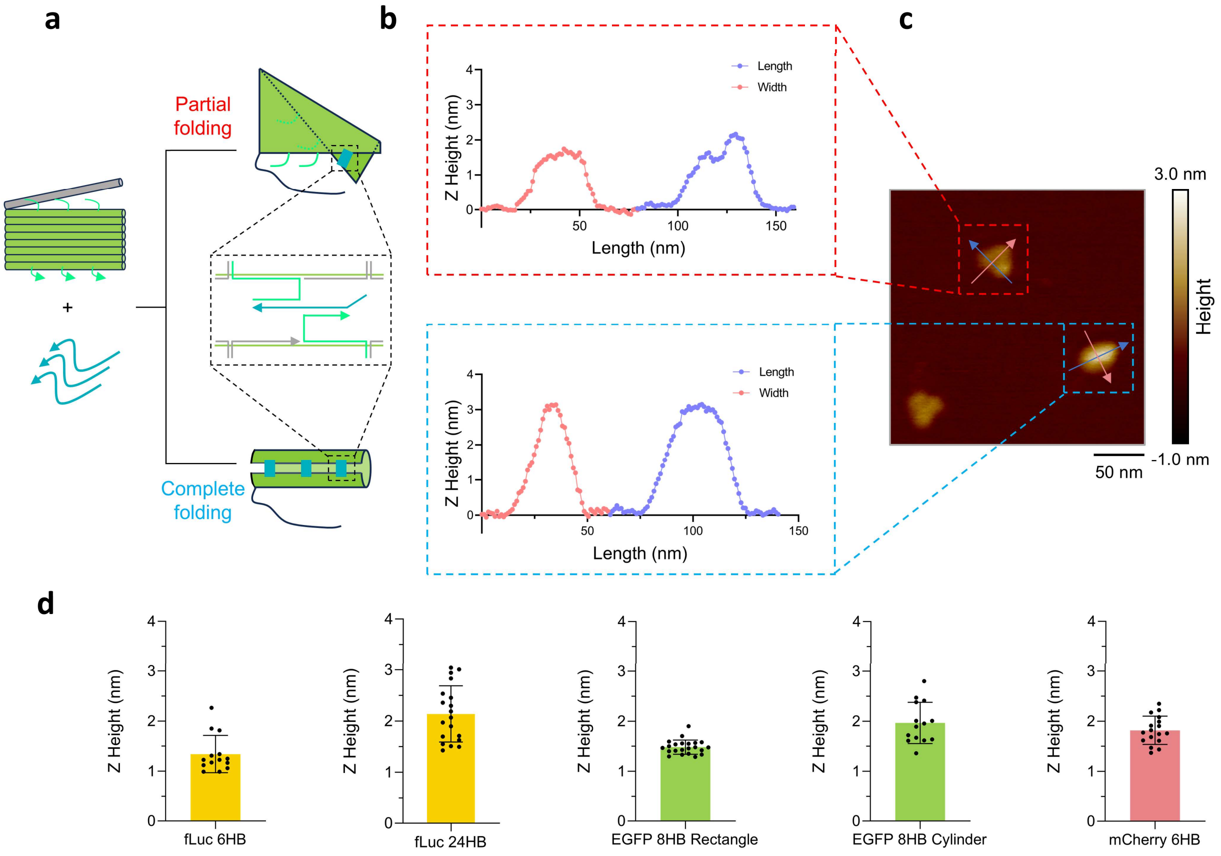

**Figure S3. Additional information regarding partial and complete folding of the EGFP 8HB cylinder and Z heights of origami.** (a) Schematic of partial folding in EGFP 8HB cylinder. Synthesis of the EGFP 8HB cylinder yielded a mixture of incompletely folded and completely folded structures, for which (b) height sections were measured from (c) 250×250 air AFM images taken using the same AFM protocol. (d) Z heights were further assessed for each origami.

### 5. Verification of AFM Distortion Using a Control DNA 6HB

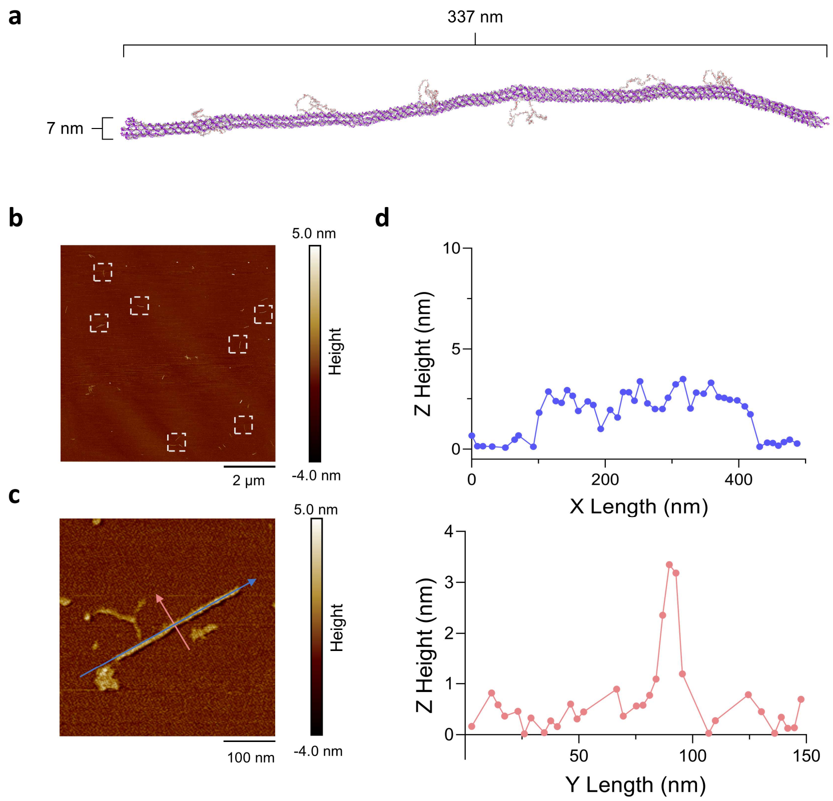

**Figure S4. Additional AFM measurements of DNA 6HB.** To validate whether our inconsistencies in AFM helical width measurements were an artefact of AFM imaging conditions, we designed (a) a DNA 6HB rod with estimated helical axial length of 337 nm and width of 7 nm. The origami was successfully synthesized, with (b) 10×10 μm air AFM reconstructions showing monodisperse structures deposited on mica, in white boxes. (c) A further magnified 500×500 nm AFM image was used for (d) height section analysis, for which a helical length of 334.7 nm and width of 15.44 nm was measured. AFM conditions were consistent with previous protocol.

### 6. Hybrid Origami Stability in 10% FBS-DMEM

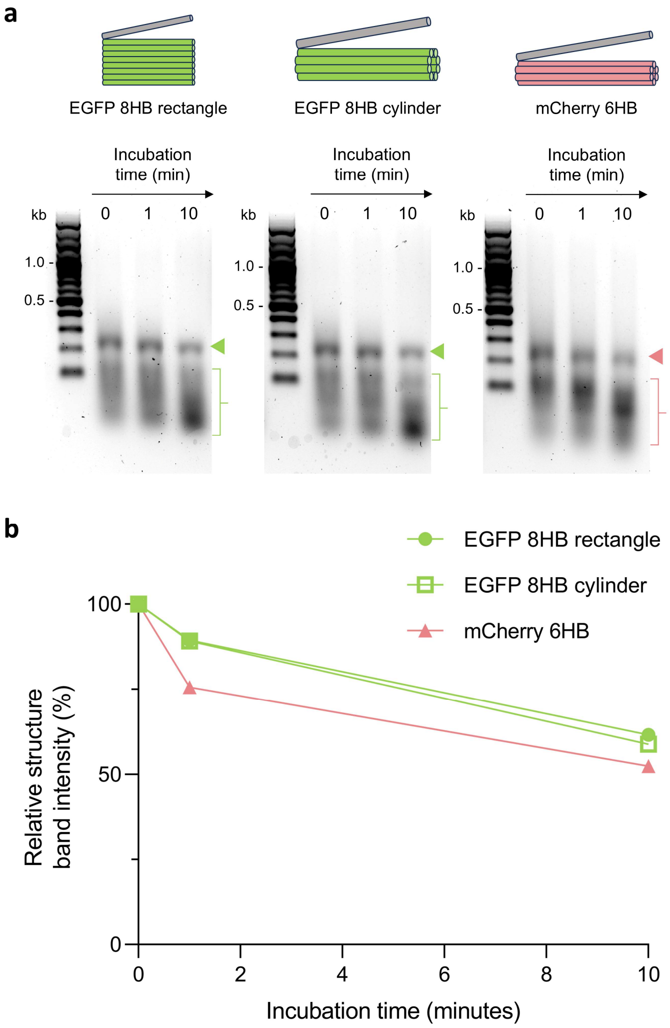

**Figure S5. Stability of hybrid origami in 10% FBS-DMEM.** (a) AGE of EGFP 8HB rectangle, EGFP 8HB cylinder, and mCherry 6HB incubated in 1× DMEM supplemented with 10% FBS for 0, 1, and 10 minutes. Triangles indicate positions of bands corresponding to origami, for which increased mobility and weakened intensity can be observed at longer incubation times. Brackets include bands corresponding to staples, for which intensity increased at longer incubation times. Together, these confirm the rapid unfolding and degradation of origami in 1× DMEM with 10% FBS. (b) Origami structure band intensity over time was measured as percentage relative to structure band intensity at initial t = 0 minutes for each structure and plotted into a scatter plot. Horizontal axis represents incubation time in 10% FBS-DMEM in units of minutes, with the vertical axis representing relative structure band intensity in units of percent. AGE conditions were consistent as previous.

### 7. Complete Agarose Gels

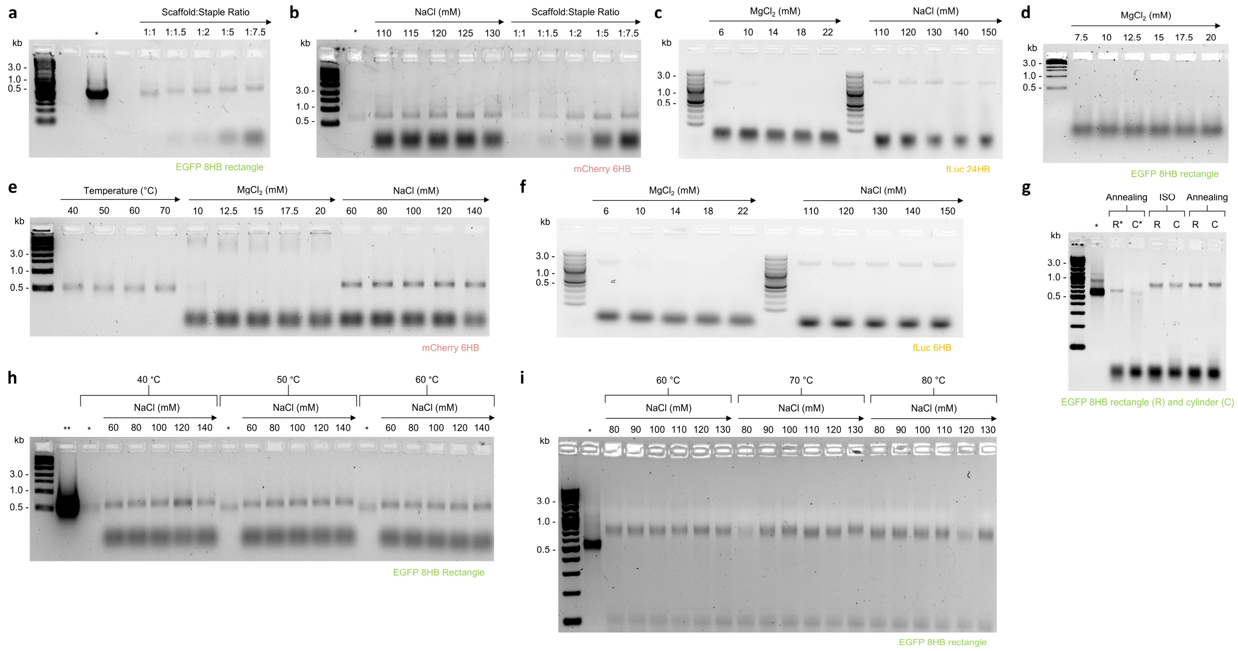

**Figure S6. Complete AGE images**. Lanes labelled with asterisks only contain only scaffold mRNA corresponding to the origami. (a), (g), (h) and (i) single asterisk lanes contain 12 µL of 20 nM EGFP mRNA, with the double asterisk lane in (i) being 12 µL 200 nM EGFP mRNA. (b) asterisk lane contains 12 µL of 10 nM mCherry mRNA. (e) 12 µL of 10 nM mCherry mRNA were incubated at initial temperatures of 40-70 °C to a final temperature of 20 °C at a ramp rate of -0.1°C/6s to assess mRNA stability. No observable difference was seen, suggesting that mRNA is stable in such protocols.

### 8. Complete Staple Sequences for Each mRNA-DNA Hybrid Origami

Table S1, S2, S3, S4, S5 correspond to staples used in fLuc 6HB, fLuc 24 HB, EGFP 8HB rectangle, EGFP 8HB cylinder, and mCherry 6HB, respectively. Position formatting is in start (ST) helix#[nucleotide position] end helix#[nucleotide position] format. Linker strands in EGFP 8HB are indicated with a cyan dot (●). Position of the mismatched 5' nucleotide is indicated in red.

**Table S1**. Staple oligo sequences for fLuc 6HB.

| Staple Sequence (5' – 3') | Position |
|---|---|
| CCGGTGCTGAGCCCTGGTAGTCGGTCTAGCTGCTCGCC | ST2[382]0[381] |
| GTCTCGGTCAGGCGGCCCCGCACGCACAGCTCCAGCTTGCC | ST3[379]5[383] |
| GGTCAGGCCCTTGGGCCCTTGTGCTGCTGTC | ST5[384]1[398] |
| CATGATGAGGCAGGGTGCCGGATGCCG | ST1[399]3[411] |
| CGGATCTTCCCAGGGGGTGTACATG | ST5[358]1[366] |
| CTCTGGACCGCTGCATGGCGCTGGTG | ST1[367]3[378] |
| GGCGGTGCCGTCCTCGGGCGTCGCCCCG | ST0[380]4[367] |
| CTGGTTCATGATCAGGTGTTCATGATCAGG | ST4[366]2[353] |
| CGTACCGCTTCATGGACCTCGTCGCTCATG | ST0[414]4[399] |
| ATCATGGCGTAGCCCCCACGCCCTTGGGCAGG | ST4[398]2[383] |
| CGCCCCGCAGCTGGTGCCGGCAGCTT | ST5[422]1[431] |
| CGTCGGTGAAGGCGATTCTTGGGGGCGTTG | ST0[447]4[432] |
| GGTCCCGGCGTTCAGGATCTTCTGCATCGATGTGGG | ST2[447]0[448] |
| GCTGGCCACGTAGTGTCCACCGGCCCTT | ST5[453]1[464] |
| GCGTAGGTGATCCACGACGCCGCTG | ST0[475]4[465] |
| CTTCTGCAGCGTGGCACGGCCTCGCCCAC | ST1[432]3[446] |
| GTGGCCTGCTTGGCCTGAACCGCACGCAGG | ST4[431]2[418] |
| CTTGCTCAGGGATGACCGCCGCTGGCGATCTC | ST1[465]3[482] |
| TGCAGCCGGGGGGCGTCTGGTTGCCGAAGATGG | ST4[464]2[448] |
| CTCCTTGCTCAAGCCGTCCTTGTCGATCACGGTGGTCACCTG | ST3[447]5[452] |
| GTCATGGTCTTGCCGGCACGCTCGGCTGATG | ST5[484]1[498] |
| CCTCGGCCAGCCTGCTCCAAGAAGTG | ST0[510]4[499] |
| CGAAGCCGTGGTGTTCAGCAGCTCCCGCTTACCGCTTCATGG | ST2[516]0[511] |
| GCGGGCAGCTCGTCAGGCCGCGTTGTA | ST5[520]1[530] |
| GGTGGTTGGTGTCCGGCGTACTTGATC | ST0[543]4[531] |
| CCCATGCTGGAAGGTCGTACTTGTCGATCAG | ST1[499]3[515] |
| CTCGTCCTTGCTCAGGGCACCACGCTCAGGATGG | ST4[498]2[481] |
| GATGTCGTTAGCCCAAAGCTGAACAGGGTGG | ST1[531]3[547] |
| AGGCTCTTGGCGAAGGGGTGGTGAACATGC | ST4[530]2[517] |
| GGTGCTCTTCAGCCGGTCCACGATGAGCACCACCACGGCG | ST3[516]5[519] |
| CGGTCCGGTGGTGATCTTCTGGATGATGGGGCACCAGGG | ST2[417]0[415] |

| | |
|---|---|
| CGGTGTCGCGAACACCACGGTGGGCTATCTCGAAGTACTCG | ST2[480]0[476] |
| GGCAGGTGGAACCCGGGGTTGTTCACGTAGCCCACGAACACCACGC | ST3[412]5[421] |
| GTGCAGGTCGTCCCAGTAGGCGATGTTCTCCTTCTCG | ST3[483]5[483] |
| GCAGGCCGGCCACGCCGTCTCGCTGCGCCGAT | ST5[548]1[563] |
| GCGGGGGGGGCGCTCGCACCACCCGGAAGC | ST4[563]2[550] |
| GAACAGGGTACATCATGGATCTTGTAGTCCTGCAGGCTCCGCAGTTTTTT | ST1[564]3[599] |
| TTTTTCATGAAGAACTGCAGGCTGTGCGTCGACCAGCTCG | ST0[589]4[564] |
| TTTTTGAACAGCTCCTCCTCGAACCGGCGCCCAGCACGGGTTTTT | ST2[598]1[589] |
| TTTTTGCAGCAGGATGCTCTAGATGTTGGGGTGCTTTTTT | ST4[591]5[591] |
| TTTTTGTCGTACTCGTGCGTCCTCCATGGTGGCTCTTA | ST1[310]0[304] |
| AATTAAGCGGCCGCTCACACGGCGATCTTGCCGCCCTGTTCTTGTGAAGCC | ST5[298]1[332] |
| GTCCACCACCACGGCGCGAAGCTCTCGGGCACGAATTTTT | ST4[333]2[310] |
| TTTTTAGGGCACCACCTTGCCCTTGGCCTCGAAGATTTTT | ST3[306]4[306] |
| GGGGCCCTTCTTGATTCTTGGTGTCCAG | ST0[347]4[334] |
| GGGGGGCATCCCGGTCCGGGCTTGTCGTCG | ST1[333]3[348] |
| GCGATGGTCTTGGGTGGCTGGTCACGAAGGTAGAAGGGGGC | ST2[352]0[348] |
| CCCTCGGGGGCGCCCAGGGTCTTGCCGGCCTTGATCAGGATCTCC | ST3[349]5[357] |
| GCACCAGCACCACCTGGTAGCCCTTGTCGTCGTCGG | ST3[548]5[547] |
| CGCAGATCAGGTGGCGGGGGCCACGGCCACACACCACGATCC | ST2[549]0[544] |
| TATTTCTTCTTACTCTTCTTTTCTCTCTTATTTCCCTTTTT | ST0[303]0[263] |
| GGGAGAGAAGAAGGGCATGGCCAGAAGGCAAGCCCCGCAGAAGGCAGCTT | ST5[248]5[297] |
| TTTTTCTTCCTACTCAGGCTTTATTCAAAGACCAAGAGGTACAGGTGCAA | ST5[198]5[247] |
| TTTTTTTTTTTTTTTTTTTTTTTTTTTTTT | ST5[168]5[197] |
| TTTTTTTTTTTTTTTTTTTTTTTTTTTTTT | ST5[138]5[167] |
| TTTTTTTTTTTTTTTTTTTTTTTTTTTTTT | ST5[108]5[137] |
| TTTTTTTTTTTTTTTTTTTTTTTTTTTTTT | ST5[78]5[107] |

**Table S2**: Staple oligo sequences for fLuc 24HB

| **Staple Sequence (5’ – 3’)** | **Position** |
|---|---|
| CAGCTTAATTAAGCGGCCGCTCACACGGCCACGCCGCCCCGCATTTTT | ST0[214]1[222] |
| TTTTTGGGCGTCCAGCTCGGTCATGGTCTTTTTT | ST1[294]2[294] |
| TTTTTGCTTCTTGGCGGTGGCGTCGGGCAGGCCGGTTTTT | ST2[222]3[222] |
| TTTTTTGCCGTGCTCCACGGCGGGGTTTTT | ST3[294]4[290] |
| TTTTTGTTCACGTAGCACCTTGCCCACTTTTT | ST7[290]8[290] |
| TTTTTCAGGGTCTTGCCGGTGGTCTCGGTCAGGCCGTAGCCCTGCCTTTTT | ST8[218]9[207] |
| TTTTTGGATGCCGGGCAGGTGGAACCGGGGCACCAGCAGGGCGCTCTGGTTTTT | ST10[207]11[207] |
| TTTTTGGCGCCGGGCTCGCCGCTGGCGATCTCGTGTTTTT | ST9[290]10[298] |
| TTTTTCAGGTTGCTCAGGTCGTACCGGAAGCCGCAGATCAGGTTTTTT | ST11[298]12[298] |
| TTTTTATCTTGTAGTCCTGCAGGCTCCTGTCGGGGATGATCTTTTT | ST12[207]13[215] |
| TTTTTAGCCCAGGGTGGGGCCACGCCCTTTTTTT | ST13[298]14[298] |
| TTTTTTGGTTGCCGAAGATGCTCTCGGGCACGATTTTT | ST14[215]15[215] |

| | |
|---|---|
| TTTTTGGGCAGGCCGGTGCTGCCGCTGCTGCTCTGGAAGCCCTGGTATTTTT | ST15[298]16[294] |
| TTTTTAGTCGTACTCGTCAGGATCTTCTGCATTTTT | ST16[215]17[211] |
| TTTTTGTCGGTCTTGCTCGCTCGTTGTAGTTTTT | ST17[294]18[294] |
| TTTTTGGCCCTTCTTGCTCAGAACTGCAGGCTGTTTTTTT | ST18[211]19[211] |
| TTTTTATGTCGTTGGCGGCCGCACGTTTTT | ST19[294]20[290] |
| TTTTTCTCGCTGCACACCACGATCCGGTCGGTGAAGGCGATGTTTTT | ST20[211]21[218] |
| TTTTTCTCATCTCGAACGGTGCCGTTTTT | ST21[290]22[287] |
| TTTTTGTGCCGGGCACCCTCCATGGTGGCTCTTA | ST22[218]23[216] |
| GGGGCGCTGTCGTCGCCCTCGGGGGTGATACCTCCTTGCTCAGG | ST10[268]10[267] |
| CTCTCCAGCTGCACCACCACGGCGGCGGGCAGTGCAGCAGGATG | ST4[268]4[267] |
| GAAGGTGTACATGTTCATGATCAGGGCGATGGTTGGCTGGTCAC | ST16[260]16[259] |
| CTCCTCGAACCGGTACATCACGTGGTGGAAGGGCAC | ST12[247]13[251] |
| CACGCTCAGGATGGCGGGCAGGAACAGCTC | ST13[250]12[246] |
| CCGGGGGGCAGGCTTGTCCCGGTCGAAGGGGTCCCGGGC | ST16[237]14[245] |
| GTGGCTGAACCGCACGCAGGGCGGGGCCCTTCTGGCTGATGC | ST14[246]18[257] |
| GGTGGGCTTGATGTTCTTGGCGTCAGGGCGTACC | ST18[241]22[244] |
| CACCACCTTTCCACCTCGATGTGGGCGTGGTTG | ST8[247]20[243] |
| GCTTCATGGCCTTGTCTCTTGGCGAAGAAG | ST22[245]11[245] |
| CTGAACAGGGTCTTGGCCACGGCCTCGCCCCAGGATGGCGCTGGTGTCCAGGTC | ST11[244]8[246] |
| CCATGCTGTTCAGCAGCTCCGTCCATGATG | ST18[258]17[268] |
| ATGATCTTTGGCCTTGATCAGGATCTCCCGGATCTTCCTTTTT | ST17[267]0[294] |
| GCCACGCCGAGTCCACGATCTCCTTCTTGCCGGTCAGGCCCTTGG | ST19[268]1[260] |
| GCACCTCGTCCACGAACACGATCTTG | ST1[259]0[249] |
| CCGCCCTTCTCTGGATGATGGGCAGCTTCTTCTGCACGTTTGAAG | ST0[250]16[236] |
| CCGTCCTTGTCGGGCCCCGCACGCACAGCTCGCCGTGCAGCCAG | ST6[248]6[247] |
| CTCGCCGGGTACTCGGC | ST22[267]21[266] |
| TTTTTTCCTCCAGGGGGTAGAAGGGGCGGTCC | ST23[287]14[271] |
| GGTGGGGCAGTGAACATGCCGAAGCGCACCACCTTGTCGATCAGGGTGGCAGCTG | ST14[272]22[266] |
| GTGTTCAGGTTCAGCCGGTCCACGTCGAAGATGTTGGGGTGCCTCGCCG | ST20[244]3[249] |
| GCGTCGTTCACCTGGCTGGCCACGTATGAACAGGGCGCCCAG | ST3[248]19[243] |
| CACGGGCATGAACGAACACCAC | ST19[242]18[240] |
| TTTTTCCACGCCGGCGATGAAGAAGTGCTCGTCCTCGTTTTTT | ST4[222]5[211] |
| TTTTTCCCAGTAGGCGATGTCGCCGCTCCGCTGGTTCACGCCTTTTT | ST6[211]7[218] |
| TTTTTGCCACCTGGTAGCCCTTGTCGTTGGTGGCCTCGGGGTTTTTTT | ST5[290]6[290] |
| GTAGGTGATGGGCCTCGAAGAA | ST21[265]8[267] |
| GGGCACCCGCTCATGATCATGGATCAGGGACTTGATCAGGCTCCCGTACC | ST8[268]20[259] |
| GCTTCATGGCCTCGGCCAGGGGCCACG | ST20[260]19[269] |
| TTTTTTTTTTTTTTTTTTTTTTTTTTTTTT | ST0[0]0[29] |
| TTTTTTTTTTTTTTTTTTTTTTTTTTTTTT | ST0[30]0[59] |
| TTTTTTTTTTTTTTTTTTTTTTTTTTTTTT | ST0[60]0[89] |
| TTTTTTTTTTTTTTTTTTTTTTTTTTTTTT | ST0[90]0[119] |
| TTTTTCTTCCTACTCAGGCTTTATTCAAAGACCAAGAGGTACAGGTG | ST0[120]0[166] |
| CAAGGGAGAGAAGAAGGGCATGGCCAGAAGGCAAGCCCCGCAGAAGG | ST0[167]0[213] |

| | |
|---|---|
| TATTTCTTCTTACTCTTCTTTTCTCTCTTATTTCCCTTTTT | ST23[215]23[175] |

**Table S3**: Staple oligo sequences for EGFP 8HB rectangle

| **Staple Sequence (5' – 3')** | **Position** |
|---|---|
| TTTTTTCTTGAAGTTCACCTTGATGTTGTGGCGGATTTTT | ST4[15]3[15] |
| TTTTTTGGACGTAGCCTTCTTAGATGGTGCGCTCCTTTTT | ST6[15]5[15] |
| GTTCTTCTCCGTCGTCCTTGAAGATTGGGCATGG | ST4[42]6[41] |
| TTTTTGTCGCCGTCCAGCTTTTTGTGGCCGTTTACTTTTT | ST8[15]7[15] |
| CGGACTTGGCCGGACACGCTGAACTTCGACCAGGATGGGCAC | ST6[42]8[49] |
| GTGGCTGTTCGAACTTCACCTCGGTTCATGTGGT | ST4[75]6[74] |
| CGGGGTAGCAGCTTGCCGTAGGTGTTCCTCGCCCTTGCTCAC | ST6[75]8[82] |
| TTTTTGTGCTCAGGTAGTCAGGGCGGACTGGTTTTT | ST3[133]2[133] |
| CAGGATGTTTCAGCTCGATGCGGTTTGCCGTAGG | ST4[108]6[107] |
| TTTTTGAAGTCGATGCCCTGCCGTCCTCCTTTTTTT | ST5[133]4[133] |
| TTTTTGCCAGGGCACGGGGTCACGAGGGTGGTTTTT | ST7[133]6[133] |
| TTTTTTTACTTGTACAGCTTTAATTAAGCGGCCGCTTTTT | ST2[15]1[15] |
| TTTTTCTTCCTACTCAGG | ST1[133]1[116] |
| TTTCTCTCTTATTTCCCTTTTT | ST8[116]8[137] |
| CCGCAGAAGGCAGCTTTTCGTCCATG | ST1[49]2[41] |
| CCGAGAGTGCACGCTGCCGTCCTCTTTTGATGCC | ST2[42]4[41] |
| GCCATGATATTGTAGTGGTCGGCGAGCTGATCCCGG | ST4[58]2[57] |
| GGAGAGAAGAAGGGCATTCGAACTCC | ST1[82]2[74] |
| AGCAGGACTGGGGGTGTTCTGCTGTTTAGACGTT | ST2[75]4[74] |
| TCTTTGCTGGTTGTCGGGCAGCAGTTTTGTGCCC | ST2[108]4[107] |
| TACTCCAGCTTCACGGGGCCGTCGCCGACATGTGATC | ST4[91]2[91] |
| GCGCTTCTTTAGAGGTACAGGTGCAAG | ST2[92]1[83] |
| CGGCGGTCATTTGGCCAGAAGGCAAGCC | ST2[58]1[50] |
| TCAGGGTGCAGCTTGCCGGTGGTGTTTTCTTCTTACTCTTCT | ST6[108]8[115] |
| CTTTATTCAAAGACCATTCGTTGGGG | ST1[115]2[107] |
| TTTTTTTTTTTTTTTTTTTT | ST0[9]0[28] |
| TTTTTTTTTTTTTTTTTTTT | ST0[49]0[68] |
| TTTTTTTTTTTTTTTTTTTT | ST0[69]0[88] |
| TTTTTTTTTTTTTTTTTTTT | ST0[89]0[108] |
| TTTTTTTTTTTTTTTTTTTT | ST0[109]0[128] |
| CACCCCGGTGAACAGCTTTGCATCGCC | ST8[50]7[59] |
| CTCGCCCTCAAGAAGTCGTGCTGCTTTTCGCGGGTCTTGTAGTTGGCTTGTCG | ST7[58]4[57] |
| CATGGTGGCTCTTATATTTCAGATGAA | ST8[83]7[92] |
| CTTCAGGGTCGGCTGAAGCACTGCACTTTCACCAGGGTGTCGCCCTGTAGTTG | ST7[91]4[90] |
| TTTTTTTTTTTTTTTTTTTT | ST0[29]0[48] |

**Table S4**. Staple oligo sequences for EGFP 8HB cylinder

| Staple Sequence (5’ – 3’) | Position |
|---|---|
| TTTTTTCTTGAAGTTCACCTTGATGTTGTGGCGGATTTTT | ST4[6]3[6] |
| TTTTTTGGACGTAGCCTTCTTAGATGGTGCGCTCCTTTTT | ST6[6]5[6] |
| GTTCTTCTCCGTCGTCCTTGAAGATTGGGCATGG | ST4[33]6[32] |
| TTTTTGTCGCCGTCCAGCTTTTTGTGGCCGTTTACTTTTT | ST8[6]7[6] |
| CGGACTTGGCCGGACACGCTGAACTTCGACCAGGATGGGCAC | ST6[33]8[40] |
| GTGGCTGTTCGAACTTCACCTCGGTTCATGTGGT | ST4[66]6[65] |
| CGGGGTAGCAGCTTGCCGTAGGTGTTCCTCGCCCTTGCTCAC | ST6[66]8[73] |
| TTTTTGTGCTCAGGTAGTCAGGGCGGACTGGTTTTT | ST3[124]2[124] |
| CAGGATGTTTCAGCTCGATGCGGTTTGCCGTAGG | ST4[99]6[98] |
| TTTTTGAAGTCGATGCCCTGCCGTCCTCCTTTTTTT | ST5[124]4[124] |
| TTTTTGCCAGGGCACGGGGTCACGAGGGTGGTTTTT | ST7[124]6[124] |
| TTTTTTTACTTGTACAGCTTTAATTAAGCGGCCGCTTTTT | ST2[6]1[6] |
| TTTTTCTTCCTACTCAGG | ST1[124]1[107] |
| TTTCTCTCTTATTTCCCTTTTT | ST8[107]8[128] |
| ●AGGCTCAATCTGATGCCAGTTAGCATATACGTGCCGT | ●ST9[91]9[127] |
| CCGCAGAAGGCAGCTTTTCGTCCATG | ST1[40]2[32] |
| CCGAGAGTGCACGCTGCCGTCCTCTTTTGATGCC | ST2[33]4[32] |
| GCCATGATATTGTAGTGGTCGGCGAGCTGATCCCGG | ST4[49]2[48] |
| GGAGAGAAGAAGGGCATTCGAACTCC | ST1[73]2[65] |
| AGCAGGACTGGGGGTGTTCTGCTGTTTAGACGTT | ST2[66]4[65] |
| TCTTTGCTGGTTGTCGGGCAGCAGTTTTGTGCCC | ST2[99]4[98] |
| TACTCCAGCTTCACGGGGCCGTCGCCGACATGTGATC | ST4[82]2[82] |
| GCGCTTCTTTAGAGGTACAGGTGCAAGCGTAGTCTAACTCGGA | ST2[83]10[58] |
| CGGCGGTCATTTGGCCAGAAGGCAAGCCGCACGGTTCACGCGCC | ST2[49]9[25] |
| ●TCCGAGTTAGACTACGGTCAATCGTATATGCACGGCA | ●ST10[58]10[94] |
| ●GGCGCGTGAACCGTGCGATTCGTCGGCCGGAGTGCCT | ●ST9[25]9[61] |
| TCAGGGTGCAGCTTGCCGGTGGTGTTTTCTTCTTACTCTTCTGCATCAGATTGAGCCT | ST6[99]9[91] |
| ACGTATATGCTAACTGCTTTATTCAAAGACCATTCGTTGGGG | ST9[122]2[98] |
| CTCCGGCCGACGAATCCACCCCGGTGAACAGCTTTGCATCGCC | ST9[56]7[50] |
| CTCGCCCTCAAGAAGTCGTGCTGCTTTTCGCGGGTCTTGTAGTTGGCTTGTCG | ST7[49]4[48] |
| TGCATATACGATTGACCATGGTGGCTCTTATATTTCAGATGAA | ST10[89]7[83] |
| CTTCAGGGTCGGCTGAAGCACTGCACTTTCACCAGGGTGTCGCCCTGTAGTTG | ST7[82]4[81] |
| TTTTTTTTTTTTTTTTTTTT | ST0[0]0[19] |
| TTTTTTTTTTTTTTTTTTTT | ST0[20]0[39] |
| TTTTTTTTTTTTTTTTTTTT | ST0[40]0[59] |
| TTTTTTTTTTTTTTTTTTTT | ST0[60]0[79] |
| TTTTTTTTTTTTTTTTTTTT | ST0[80]0[99] |
| TTTTTTTTTTTTTTTTTTTT | ST0[100]0[119] |

**Table S5**. Staple oligo sequences for mCherry 6HB

| Strand Sequence (5' – 3') | Position |
|---|---|
| CTGGGTATCTCGAACTCTGAACCGCATGAACTCCTTTTTT | ST0[32]2[7] |
| AGGTAGTCTCAGGAACCTCGGCGTCGTAGTGGCC | ST2[80]1[104] |
| TGCACCTGTGGCCGTTCACGCTAGGCCTTGCTGC | ST2[36]1[60] |
| TGCTCACCATTTCATCACGGCGGGGTGCTTCACGTGCCCTCCATG | ST3[42]2[37] |
| GGCGCCGGCCGCTGCTTGATCTGGCTGGTGATGTCCTTTTT | ST2[124]1[155] |
| TGGGGAAGTTCTCGTACTGCTCCACTTTTT | ST3[130]3[159] |
| TTTTTGATGGTGTAGTCCTCGTTGTCGCCCTTCAG | ST2[159]2[125] |
| GCCCTCGGGGAAGCCTCCCAGCCCATGGTACATCCGCTCGCTGCTGGCTCAGCTTC | ST3[75]2[81] |
| TTTTTTTTTTTTTTTTTTTT | ST6[6]6[25] |
| TTTTTAGCTTGATGTTCACGTTGTAGGCGCCCGGTGCTGTGCCGGCCCTTTTT | ST0[155]5[152] |
| CGTACAGGCAGGGGGCCGCCCT | ST1[61]0[55] |
| TGGTCACAGAAGGCAAGCCCCGCAGAAGACGGTCACCACGTGAA | ST0[54]3[74] |
| GCGAAGTGAACTGGGGGCGGGGATGTCCCGCTCCCACTCCGCC | ST0[76]4[62] |
| TTTTTTCGGCCCGGGTGCCCCGCAGCTTCACGTCCATGCCGCGGGC | ST4[152]0[121] |
| AGCTGCCTTCAGCTTCATCCTCGGGGTCTTCTTCTGCAACTCGCCGTCCTGCAG | ST0[120]4[95] |
| GCCGTCACGGGCTTCTTGGCCTTGTAGGCAGTTGCCGCTCTTG | ST1[105]5[111] |
| TACAGCTCCTTGTAGATGATCACGGGGCCGTCGC | ST5[112]3[129] |
| TTTTTTCGCCCTCGCCCTCGGCCCTCGTAGGGTACAGGTGCATTCTTA | ST1[3]4[29] |
| CTCTTCTGTCCTCCTCGCCCT | ST4[28]3[41] |
| GTCCTCGAAGGGTGGCTCTTATATTTCAGGGAGAGAAGAAGG | ST4[61]5[45] |
| GCATGGCCCTTCAGCTTGGCGGT | ST5[46]0[33] |
| GCTGCTGTCCTGGGTCGCAGCTTAATTAAGCGGCCGCTTGGTCTTCTGTCCCAG | ST4[94]0[77] |
| TTTTTTGATGATGGCCATGTTTTTCTCTCTTATTTCCCTTTTT | ST3[7]4[0] |
| TTCAAAGACCAAGAGGCCGGCCCTTTTT | ST5[5]0[3] |
| TTTTTTTTTTTTTTTTTTTT | ST6[66]6[85] |
| TTTTTTTTTTTTTTTTTTTT | ST6[46]6[65] |
| TTTTTTTTTTTTTTTTTTTT | ST6[106]6[125] |
| TTTTTTTTTTTTTTTTTTTT | ST6[86]6[105] |
| CTTCCTACTCAGGCTTTA | ST6[126]6[143] |
| TTTTTTTTTTTTTTTTTTTT | ST6[26]6[45] |

#### 9. Thermal Incubation Protocol Details

**Table S6.** Specific thermal annealing protocols

| Protocol | Temperature gradient (°C/min) | Initial temperature (°C) | Final Temperature (°C) | Time (hours) |
|---|---|---|---|---|
| Standard Annealing | -0.146 | 55 | 20 | 4 |
| Isothermal 55 °C | -0.000 | 55 | 20 | 4 |
| $T_{max}$ = 40°C | -0.083 | 40 | 20 | 4 |
| $T_{max}$ = 50°C | -0.125 | 50 | 20 | 4 |
| $T_{max}$ = 60°C | -0.167 | 60 | 20 | 4 |
| $T_{max}$ = 70°C | -0.208 | 70 | 20 | 4 |
| $T_{max}$ = 80°C | -0.250 | 80 | 20 | 4 |

#### 10. Complete Staple Sequences for Control DNA 6HB Origami

**Table S7.** Staple oligo sequences for DNA 6HB origami

| Staple Sequence (5' – 3') | Position |
|---|---|
| AAGAATATACAAATCACCAGAGCCGCCGCGATTGGTGGCGAGAAAGGAAGGGAATTTTT | ST2[55]0[25] |
| GCTAGGGGCGAACGCCTTGATATTCACAGAACCACTCTTACC | ST1[49]3[62] |
| GCCACCAAACAAATGACGGGGAAAGCCGCGCTGGCGTTAAAT | ST4[69]2[56] |
| AGTATAAATAAGGCAAGTGTAGCGGTCAAGAGCTTAAATCCT | ST3[63]5[76] |
| CCGATTTCGCTGCGCGACCGTGTGATAAAGCCAACCAGAGCC | ST0[83]4[70] |
| ATGGTTTCTTAATTCACCCTCAGAGCCAGCCAGAAGGAGCCC | ST2[97]0[84] |
| ACCACACCCTAAAGTGGAAAGCGCAGTCGAACCGCGAGAATC | ST1[91]3[104] |
| ACCCTCATCTGAATCACTAAATCGGAACCCGCCGCAAATTTA | ST4[111]2[98] |
| GCCATATCTGACCTGCTTAATGCGCCGCCGTAAAGTTACCGT | ST3[105]5[118] |
| GAGGTGCTACAGGGGTTAATTTCATCTTTTAACAAAGCCGCC | ST0[125]4[112] |
| TCAAATATTAGGCACACCGGAACCGCCTAGCGTCATGGGGTC | ST2[139]0[126] |
| TATGGTTAGTTTTTTACATGGCTTTTGAGAGCCACGAGGCAT | ST1[133]3[146] |
| GGAACCATGATACACATCACCCAAATCAGCTTTGAAACTTTT | ST4[153]2[140] |
| TTTCGAGGCGAGAACGAGCACGTATAACCGTGAACGGAGTGT | ST3[147]5[160] |
| CCCACTAGTGCTTTGCAAGACAAAGAACCCAGTAAAATCACC | ST0[167]4[154] |
| ATGCAAAGTACCGATTGCCATCTTTTCAATAAGTTGCGATGG | ST2[181]0[168] |
| AGAATCATATCAGGTTAACGGGGTCAGTTTAGCGTCAAAAGG | ST1[175]3[188] |
| CCCCTTAGCCTTGAAGACGGGAACCGTCGAGCGGGAATGCTG | ST4[195]2[182] |
| TAAAGTAATATGTAAGCTAAACAGGAGGACCAGTGGTAACAG | ST3[189]5[202] |
| TCTTTTCCCGATTATGGGTTATATAACTATTCTGTTCATAGC | ST0[209]4[196] |
| CCTCCGGTAAACAACGTTTTCATCGGCAATAAACATGGTTTT | ST2[223]0[210] |
| TTTAGACGCCAGGGGTTAATGCCCCCTGTGTAGCGCATGTTC | ST1[217]3[230] |
| GTCAGACCCTATTTTTTGCGTATTGGGCAGGAACGTTTTTAA | ST4[237]2[224] |
| AGCTAATGACTACCGTACGCCAGAATCCGAGGCGGCGGAACC | ST3[231]5[244] |
| GCGGGGATGAGAAGTCATAGGTCTGAGAGCAGAACCTTTAGC | ST0[251]4[238] |
| TGAATTTACAATAGTAGCGACAGAATCACTGAAACCCAACGC | ST2[265]0[252] |
| ATAATCAGAATCGGATGAAAGTATTAAGTAATCAGATAAGTC | ST1[259]3[272] |
| AGCACCGAGGCTGACCAGCTGCATTAATGTGAGGCTCAATAG | ST4[279]2[266] |
| CTGAACAAGAAGAGCACCGAGTAAAAGATGTCGTGGACTCCT | ST3[273]5[286] |
| GGAAACCGTCTGTCAGATTAAGACGCTGAGAAAAACGATAGC | ST0[293]4[280] |
| CATAGCGAATTTACAAACGTCACCAATGAGGATTACCAGTCG | ST2[307]0[294] |
| CAAATTACCGCTTTGGATTAGCGGGGTTAGGCCGGGAGCATG | ST1[301]3[314] |
| ATTAGCATTGCTCAGTTGCGCTCACTGCACCGTTGTTGAAAA | ST4[321]2[308] |
| TAGAAACAGAATCCTAGCAATAGATAGATAATTGCGTACCAG | ST3[315]5[328] |

| | |
|---|---|
| CTCACATACCCTTCATTAATTTTCCCTTCAATCAACATTACC | ST0[335]4[322] |
| AATCGTCTCCTTATCAGCAAAATCACCAAGTGCCGGAGCTAA | ST2[349]0[336] |
| AAAGCGTAATGAGTTCGAGAGGGTTGATTTAGAGCCATTCCA | ST1[343]3[356] |
| TTGGGAAATAAGTAAGCCTGGGGTGCCTAAGAATATTCTGTA | ST4[363]2[350] |
| AGAACGGACCTTGCCGTGGCACAGACAAAGTGTAATAGCCCG | ST3[357]5[370] |
| AGCATAATATTTTTTATGTGAGTGAATAGTATTAAGAGCCAT | ST0[377]4[364] |
| TACATAATCATCGAAATTATCACCGTCATGTATCAAGCCGGA | ST2[391]0[378] |
| TATTAGTACATACGCCGTACTCAGGAGGAAAGGTGGAACAAG | ST1[385]3[398] |
| ATTCATTTTTAGTACACAATTCCACACACTTTAATGAAACAG | ST4[405]2[392] |
| CAAGCCGTTTAATGGCGCGAACTGATAGATCCGCTCCGCCAC | ST3[399]5[412] |
| AATTGTTCCCTAAATTTGAATTACCTTTTTTTTATGGAAATT | ST0[419]4[406] |
| ATTTAACTCATTACGGAGGGAAGGTAAAACCGCCATGTGTGA | ST2[433]0[420] |
| CATTAAATGTTTCCCCCTCAGAACCGCCGATTGAGCGCGCCC | ST1[427]3[440] |
| TTCAACCACCCTCAATCATGGTCATAGCAATACCGTAATTAC | ST4[447]2[434] |
| AATAGCAACAAAATAACGAACCACCAGCATTCGTAGAGCCAC | ST3[441]5[454] |
| AGCTCGAAGAAGATAGGGTTACAAGAAAAGCAAATGGCGACA | ST0[461]4[448] |
| TTCTGAATTATCCGTACCAGCGCCAAAGATTTTCAGGTACCG | ST2[475]0[462] |
| AGGTGAGATCCCCGGGGATAGCAAGCCCTATGGTTGTATTCT | ST1[469]3[482] |
| AAATTCAAATAGGAGTCGACTCTAGAGGGCGGTCAATTATAC | ST4[489]2[476] |
| AAGAACGTGTTTGGGTATTAACACCGCCCCTGCAGACCCATG | ST3[483]5[496] |
| TTGCATGTGCAACAAATATAATCCTGATCGAGGCGCAATAGA | ST0[503]4[490] |
| GATGGCACCGACTTATAAGTTTATTTTGACACTGAGCCAAGC | ST2[517]0[504] |
| GCTGAGAGGCCAGTGTTTCGTCACCAGTCCACGGAGCGGGAG | ST1[511]3[524] |
| AAAGACAACAAACTCGTTGTAAAACGACGCCAGCAATCAGAT | ST4[531]2[518] |
| GTTTTGACCTGATTGCAAATGAAAAATCGTCACGAACAACGC | ST3[525]5[538] |
| TTTCCCATAAAGCAATTATCATCATATTAGCCTTAGAAACGC | ST0[545]4[532] |
| CCAGAAGGCTATTTTAAAGGTGGCAACAATTCCACCCAGGGT | ST2[559]0[546] |
| GCTGAACGGTAACGAGACAGCCCTCATAACATACATGCACCC | ST1[553]3[566] |
| AGAAAATGTTAGCGAGGCGATTAAGTTGCTCAAATGAAACCA | ST4[573]2[560] |
| AGCTACAGAACAAAATCAAACCCTCAATTGCTGCATAACGAT | ST3[567]5[580] |
| GGGGATGCAATATCATTATCATTTTGCGATTTTATCAAACGT | ST0[587]4[574] |
| AAAGTTTAGAGATACCTTATTACGCAGTTTTGTCGGCGAAAG | ST2[601]0[588] |
| TTGGCAACCAGCTGTCTTTCCAGACGTTTAAGACTACCCACA | ST1[595]3[608] |
| GCATGATAGTAAATTCTTCGCTATTACGATCAACAAATTTTA | ST4[615]2[602] |
| AGAATTGCGTTATTGTTGAAAGGAATTGGCGGGCCGAATTTT | ST3[609]5[622] |
| GATCGGTAGGAAGGATCCTTTGCCCGAAAGTTAAGAGAACTG | ST0[629]4[616] |
| ACAACTCCAAGAAAAATAATAACGGAATGGATTTTGAAGGGC | ST2[643]0[630] |
| AAATATCCTGTTGGGCTAAACAACTTTCGAAACGCCAATGAA | ST1[637]3[650] |

| | |
|---|---|
| AACCGAGAACAGTTTTCAGGCTGCGCAATTTAGGACAATTCG | ST4[657]2[644] |
| ATAGCAATTACAAAGCACTAACAACTAATTCGCCATCAGCGG | ST3[651]5[664] |
| AGCGCCATAGATTAAGAAGTATTAGACTTAGCTATAGAAGGA | ST0[671]4[658] |
| CAATACTGCTAAGCAGATAGCCGAACAAATAGAAACAGGCAA | ST4[699]0[672] |
| TGCCGGAAACGGAACAACTAAAGGAATTGCGACCGCTTCTGG | ST0[688]0[689] |
| CAATAGATACCTTTATTTCAACGCAAGGTTCCGGCAATAATA | ST1[679]5[706] |
| GCCAGCTATAAAAAACCCTGTAATACTTATAAATATAGCGTC | ST0[713]4[700] |
| CCAAAAACTCAAATTAGTAAAATGTTTACACGTTGCACTCCA | ST2[727]0[714] |
| AACCCTCAAGATCGAAAATCTCCAAAAAGGGGTAAGCTTTAA | ST1[721]3[734] |
| GCCAGAGAAAGGCTGTATCGGCCTCAGGATATATTGGTTGTA | ST4[741]2[728] |
| ACAGTTCCTAAATCTTAAATGCAATGCCGACGACACCAAAAG | ST3[735]5[748] |
| AGGGGACTGAGTAACTCAGAGCATAAAGAGAAAACAAGTTTT | ST0[755]4[742] |
| TTAAGCAATCAAAAGAACTGGGGCTTTTTAATTGTCAGTTTG | ST2[769]0[756] |
| GTAAAGACATCTGCATCGGTTTATCAGCATTTTAAATCAGGT | ST1[763]3[776] |
| TTATGCGTTGCTTTGCATCGTAACCGTGTTCAAAAAGCAAAA | ST4[783]2[770] |
| CTTTACCAAGAATTGGGTGAGAAAGGCCGATGGGCCGAGGTG | ST3[777]5[790] |
| TGGTGTAGGAGACAATACAGGCAAGGCACTGACTAAATTACC | ST0[797]4[784] |
| ACATCCAAAAGCGGATTTCAACTTTAATTAAACAGGTCACGT | ST2[811]0[798] |
| CACCATCGGGATAGCTTGATACCGATAGTGGTTTAATTGCAT | ST1[805]3[818] |
| CTTGAGATTGCGCCGGATTGACCGTAATAATATGAAGCATTA | ST4[825]2[812] |
| CAAAAAGTAGTAGTTATTCAACCGTTCTAAACGGCGACAATG | ST3[819]5[832] |
| TGGGAACAGCTGATATCAATTCTACTAAATTAAGAAATTGGG | ST0[839]4[826] |
| GCTGAAACTTCAAAGAAACACCAGAACGCCATCGCTTCTCCG | ST2[853]0[840] |
| TGCCGGACGTCGGACCACGCATAACCGACTGACGATATCGCG | ST1[847]3[860] |
| GCTTGCCTATATTCAGCGAGTAACAACCGAGGGTAGGCGCGA | ST4[867]2[854] |
| TTTTAATCATTTGGGCTATTTTTGAGAGAAATGTGGGTCGCT | ST3[861]5[874] |
| CAACATTATCTACATTTAGCTATATTTTTCGAGCTGAATAAG | ST0[881]4[868] |
| TGGTCAAACCGGAATAACAAAGCTGCTCGCAGGGACTTTCAT | ST2[895]0[882] |
| TCAGGTCTAGCCAGGTTAAAGCGAAACAATCAACGGCAAACT | ST1[889]3[902] |
| TACCCAAAAGTACAGTCTGGCCTTCCTGATTGCCTTCGCAAA | ST4[909]2[896] |
| CCAACAGATACATTGAGAGTCTGGAGCAAATTCGCACGGAGA | ST3[903]5[916] |
| CAAAAATAACAAGAAGTTTGACCATTAGGTCAGGATATTCAT | ST0[923]4[910] |
| GAACGAGTAATTGCGTAATCTTGACAAGCATCGCCACGCCAT | ST2[937]0[924] |
| TGAACGGAATAGGATGATAAATTGTGTCATCAAGATCCTTTT | ST1[931]3[944] |
| GACCTTCGAAATCCCTCATTTTTTAACCTAATCGTATTCTGC | ST4[951]2[938] |
| GATAAGAATTCCCAAAAACTAGCATGTCAAATCAGGCGACCT | ST3[945]5[958] |
| TTTTGTTAATCATACCATATAACAGTTGGGTCATTGCTGGCT | ST0[965]4[952] |
| CTGGAAGAGAGCTTGTGTACAGACCAGGGTTACTTATTAAAT | ST2[979]0[966] |

| | |
|---|---|
| CGGTTGAAATTCGCAGCCGGAACGAGGCATGAACGAATTGCT | ST1[973]3[986] |
| AGGACAGGCAGACGTAATATTTTGTTAATAATCAGACGGTGT | ST4[993]2[980] |
| GAATATACTAAAGTAAAAGCCCCAAAAATAAACGTGTCAATCATAAGGGAACCGTTTTT | ST3[987]5[1017] |
| TTTTTGCAAATATTTAAATTGCAGGAAGATTGTATAATTTTT | ST0[1021]1[1021] |
| TTTTTTTTTTGAAAGCGAAAGGGAATCATAATTACTAGAAATTTTT | ST1[20]2[18] |
| TTTAAGCCTGTTTAGTATCATATGCGTTAAACACCGAGCGGGC | ST3[20]1[48] |
| CATTAAACCACCCTGCTCAACAGTAGGGGAAATACCGTAACC | ST5[77]1[90] |
| TCCAGTACCCTCAGCGCCAACATGTAATTATTTTACGCGTAC | ST5[119]1[132] |
| ACTGGTATAATCAATAAGAGAATATAAATCCAATCCCTCGTT | ST5[161]1[174] |
| TGCCCGTTTTTCGGCCAGACGACGACAACTTAGGTAAGGGAT | ST5[203]1[216] |
| TATTATTAGTTTGCGCGCCTGTTTATCAATCAAAATGTTTTT | ST5[245]1[258] |
| CAAGAGAAAACCATTAATATCCCATCCTATAGCTTCATCACG | ST5[287]1[300] |
| GCGGATAGTAGCACTAATCGGCTGTCTTGCTATTATGACCTG | ST5[329]1[342] |
| GAATAGGCCGACTTACCAAGTACCGCACATCAATAGAATGGC | ST5[371]1[384] |
| CCTCAGATATTGACTTTCATCGTAGGAAAATTTCAACATCGC | ST5[413]1[426] |
| CACCCTCACAAAAGCAGATATAGAAGGCTAATGGAAAAACAG | ST5[455]1[468] |
| TACCGTATCACAATTTTTAGCGAACCTCATTCATCGTGCCAC | ST5[497]1[510] |
| CTGTAGCTATAAAAAATCAAGATTAGTTGAGCGGATCACCTT | ST5[539]1[552] |
| CTAAAGTATGTTAGCCTGAATATATCAGGAGTAACTGGTCAG | ST5[581]1[594] |
| CTGTATGACCCAAACCCAATAATAAGAGGTATTAATTATCTA | ST5[623]1[636] |
| AGTGAGAAGTTACCCTTACCGAAGCCCTAGGATTTGAGCCGT | ST5[665]1[678] |
| ATTTTTTGACTGGATTCATTGAATCCCCCATTATGTTTTTAG | ST5[707]1[720] |
| GAGCCTTGCAAAAGGAGAATGACCATAAATAAAGCTGTGTAG | ST5[749]1[762] |
| AATTTCTCATTGTGTTATAGTCAGAAGCATAAATCGTCAAAT | ST5[791]1[804] |
| ACAACAAAGTAGTAGGAAGCCCGAAAGAAGGTGGCAAATTAA | ST5[833]1[846] |
| GAGGCTTATTCAGTTCAAAGCGAACCAGTAACCTGAAGGCTA | ST5[875]1[888] |
| TTTGTATAACCGGATTAGAGAGTACCTTTAGATTTGAATCGA | ST5[917]1[930] |
| GCTCCATCGCATAGTTTGCGGATGGCTTTTTCATTTGTACCC | ST5[959]1[972] |
| GAGAAGATACATTTGTTTTAAGAAAAGGGAATCGTCTTGCGG | ST2[693]2[694] |
| TTTTTTGTTTTAAATATGCAAATGCTGTAGCTCAACACTCTACCATCCACACACTAC | ST2[1014]3[1029] |
| TTTTTTGTTGAGGCAGGTCAGACCAGCATTGACAGGAGTTTTT | ST5[20]4[21] |
| TTTTTAACTGACCAACTTTGAAAG | ST4[1017]4[994] |

### 11. Additional SNUPI Simulation Parameters

**Table S8.** Average intrinsic properties of RNA-DNA hybrids obtained by SNUPI molecular dynamics simulations

| Properties | Unit | Regular | Nicked | CO-nick | Double CO | Single CO |
|---|---|---|---|---|---|---|
| $\Delta_{x1}$ | nm | -0.1682 | -0.1650 | -0.1664 | -0.8971 | -0.8995 |
| $\Delta_{x2}$ | nm | 0.1682 | 0.1650 | 0.1664 | 0.8971 | 0.8995 |
| $\Delta_{y1}$ | nm | 0.0555 | 0.0535 | 0.0624 | -0.2171 | -0.2079 |
| $\Delta_{y2}$ | nm | -0.0555 | -0.0535 | -0.0624 | 0.2171 | 0.2079 |
| $\Delta_{z1}$ | nm | -0.0035 | 0.0203 | 0.0288 | -0.0658 | -0.0388 |
| $\Delta_{z2}$ | nm | 0.0035 | -0.0203 | -0.0288 | 0.0658 | 0.0388 |
| $\Theta_{x1}$ | deg | -15.2614 | -14.1432 | -13.7388 | -10.4948 | -15.0839 |
| $\Theta_{x2}$ | deg | 15.2614 | 14.1432 | 13.7388 | 9.1807 | 13.6431 |
| $\Theta_{y1}$ | deg | -4.3730 | -3.1797 | -4.3317 | 1.7553 | -2.8749 |
| $\Theta_{y2}$ | deg | 4.3730 | 3.1797 | 4.3317 | -1.2373 | -1.9552 |
| $\Theta_{z1}$ | deg | 0.0293 | -0.2791 | 0.8487 | -34.2206 | -38.9939 |
| $\Theta_{z2}$ | deg | -0.0293 | 0.2791 | -0.8487 | 7.0168 | 12.9907 |
| $EA$ | pN | 1874.1320 | 1481.1472 | 1551.0965 | 2987.5783 | 2270.1834 |
| $GA_y$ | pN | 781.4645 | 504.8950 | 591.4058 | 1050.3941 | 889.1491 |
| $GA_z$ | pN | 399.6779 | 306.5581 | 340.8309 | 568.0062 | 442.9013 |
| $GJ$ | pN nm$^2$ | 319.4592 | 133.6630 | 170.4630 | 234.9423 | 191.5606 |
| $EI_y$ | pN nm$^2$ | 155.6086 | 124.2667 | 128.4138 | 322.9594 | 276.5894 |
| $EI_z$ | pN nm$^2$ | 250.4547 | 195.2718 | 202.5125 | 269.8093 | 221.3445 |

| | | | | | | |
|---|---|---|---|---|---|---|
| $g(\Theta_x,\Theta_y)$ | pN nm$^2$ | 23.6679 | 2.5161 | 5.3306 | -13.1454 | -15.6757 |
| $g(\Theta_z,\Theta_z)$ | pN nm$^2$ | -0.4919 | 12.2400 | 18.5200 | -12.1391 | -24.9004 |
| $g(\Theta_y,\Theta_z)$ | pN nm$^2$ | 3.5975 | 2.8587 | 1.9402 | -0.1586 | -7.7172 |
| $g(\Delta_x,\Delta_y)$ | pN | 253.9537 | 160.8202 | 121.8705 | 894.1666 | 679.0053 |
| $g(\Delta_x,\Delta_z)$ | pN | 4.3471 | 21.4768 | 30.2929 | 383.6235 | 259.5557 |
| $g(\Delta_y,\Delta_z)$ | pN | 3.1783 | -7.0446 | 12.8608 | 185.7955 | 116.3319 |
| $g(\Delta_x, \Theta_x)$ | pN nm | -219.0912 | -100.4597 | -96.0159 | -3.5904 | 8.6364 |
| $g(\Delta_x, \Theta_y)$ | pN nm | -128.5540 | -63.4921 | -96.2464 | -6.9514 | 10.5588 |
| $g(\Delta_x, \Theta_z)$ | pN nm | -9.4942 | -40.2170 | -10.8294 | -24.0296 | 132.9342 |
| $g(\Delta_y, \Theta_x)$ | pN nm | -109.3867 | -55.3284 | -57.1980 | 42.7305 | 24.1726 |
| $g(\Delta_y, \Theta_y)$ | pN nm | 5.8826 | 22.7348 | 14.9791 | -4.8358 | 8.0022 |
| $g(\Delta_y, \Theta_z)$ | pN nm | -0.4455 | -25.3409 | -43.9547 | 14.2834 | 48.7751 |
| $g(\Delta_z, \Theta_x)$ | pN nm | -3.2659 | -100.3379 | -88.3834 | 95.7297 | 88.1513 |
| $g(\Delta_z, \Theta_y)$ | pN nm | -1.8798 | -11.7841 | -21.3170 | 18.5331 | -4.4003 |
| $g(\Delta_z, \Theta_z)$ | pN nm | -38.0305 | -48.0179 | -41.4560 | -29.2270 | 5.0872 |

**Table S9.** Main parameters of SNUPI finite element analysis

| Options | Abbreviation | Value | Unit |
|---|---|---|---|
| **1. Base-pair (BP) and crossover (CO) steps** | | | |
| - Order of coefficient function | BP_CF_IND | 2 | |
| **2. Single-stranded DNA (ssDNA)** | | | |
| - Contour length per nucleotide for short ssDNA | SSD_LCT1_S | 0.38 | [nm/nt] |
| - Contour length per nucleotide for long ssDNA | SSD_LCT1_L | 0.68 | [nm/nt] |
| - Persistence length for long ssDNA | SSD_LPB_L | 0.67 | [nm] |
| - Stretching rigidity when stretched | SSD_EA_H | 710 | [pN] |
| - Stretching rigidity when relaxed | SSD_EA_L | 15 | [pN] |
| - Order of coefficient function | SSD_CF_IND | 3 | |
| **3. Single-stranded RNA (ssRNA)** | | | |
| - Contour length per nucleotide for short ssRNA | SSR_LCT1_L | 0.383 | [nm/nt] |
| - Contour length per nucleotide for long ssRNA | SSR_LCT1_L | 0.606 | [nm/nt] |
| - Persistence length for long ssRNA | SSR_LPB_L | 0.606 | [nm] |
| - Stretching rigidity when stretched | SSR_EA_H | 1512 | [pN] |
| - Stretching rigidity when relaxed | SSR_EA_L | 8 | [pN] |
| **4. Electrostatic Interaction** | | | |
| - Temperature | ES_TEMP | 300 | [K] |
| - NaCl concentration | ES_NaCl | 300 | [mM] |
| - Effective charge per base pair | ES_QEFF | 1 | [e] |
| - Cutoff distance | ES_R_CUT | 2.5 | [nm] |
| - Order of coefficient function | ES_CF_IND | 1 | |
